# Single cell transcriptome analysis of decidua macrophages in normal and recurrent spontaneous abortion patients

**DOI:** 10.1101/2021.03.23.436615

**Authors:** Qingliang Zheng, XiangHong Xu, Fenglian Yang, Haili Gan, Yongbo Zhao, Xiaoping Wan, Liping Jin

## Abstract

Due to the heterogeneity and different polarization state of decidual macrophages (dMΦ), they play an important role during the pregnancy, but their definition and exact function remain elusive. We isolated CD14^+^CD45^+^ dMΦ from the normal or RSA decidua by flow cytometry, followed by single cell RNA sequencing (scRNA-seq). In total, 23,062 single-cell transcriptomes of macrophage were profiled (12,470 Normal and 10,592 RSA), which were divided into 13 major clusters via T-distributed stochastic neighbor embedding (t-SNE) visualization. We observed that there is higher percentage composition of M1 cells (70.6%) in the normal decidua, and higher percentage composition of M2 cells (68.3%) in the RSA decidua. We identified new markers (M1: S100A8, S100A9, M2: SELENOP, FOLR2, RNASE1) and secreted cytokines (M1: IL1β, TNFSF13B and MMP9; M2: CCL3, CCL4) of the dMΦ. We found that cluster 10 as the specific cluster of dMΦ highly expressing BAG3 in normal group and cluster 7 specific highly expressing CXCL9/10/11 chemokine. After pseudo-time trajectory analysis, we found that the dMΦ formed a continuous “V-shaped” trajectory, with M1 and M2 type cells mainly occupying the two heads. We found that NFκB1, MYC and TCF12 acted as the key transcription factors of dMΦ. Our study redefined the polarization state and physiological characteristic of dMΦ in early normal and RSA pregnancy, which suggests a novel view and therapeutic target for spontaneous abortion prevention.

## INTRODUCTION

The microenvironment of normal pregnancy requires dynamics precise regulation of various immune cells during the pregnancy (Erlebacher, 2013). If external stimulation or physiological factors disrupt this precisely regulation, this will lead to pregnancy complications (Perez-Sepulveda, Torres, Khoury, & Illanes, 2014). There are many kinds of immune cells at the maternal-fetal interface. Among them, decidual macrophages (dMΦ) are the second largest group of cells and comprise 20– 30% of all leukocytes at the maternal-fetal interface (Gomez-Lopez, StLouis, Lehr, Sanchez-Rodriguez, & Arenas-Hernandez, 2014). Because of this special immune microenvironment, there are many differences in function and characteristics between dMΦ and macrophages from peripheral blood. However, the surface marker genes of macrophages from peripheral blood (such as CD209, CD80) were still used as FACS markers to isolate dMΦ (Svensson et al., 2011). It is necessary to explore the specific new marker of dMΦ using new methods. At present, several research groups have reported the composition of various cell types including dMΦ in the whole maternal-fetal interface (Tsang et al., 2017; Vento-Tormo et al., 2018), but the precise markers of dMΦ under this specific condition need to be furtherly illustrated.

Macrophages display important roles in the pregnancy process via their polarization, through which macrophages can differentiate into specific phenotypes and have specific biological functions in response to microenvironmental stimuli (Aagaard et al., 2014; Raj, Bonney, & Phillippe, 2014). Macrophages can be polarized into classically activated (M1) and alternatively activated (M2) macrophages based on their activation states (Houser, Tilburgs, Hill, Nicotra, & Strominger, 2011). Actually, M1 macrophage are potent effector cells involved in Type 1 T helper (Th1) responses, such as cytotoxicity toward microorganisms and enhanced production of pro-inflammatory cytokines. In contrast, M2 macrophages suppress the inflammatory response, skew the immune response toward Th2-associated immunity, promote tissue remodeling and induce angiogenesis (Svensson-Arvelund & Ernerudh, 2015). Macrophage polarization is crucial for tissue repairing and homeostasis maintenance (PrabhuDas et al., 2015; Tilburgs, Evans, Crespo, & Strominger, 2015). The state of macrophage activation is related to many kinds of pregnancy complications, during this specific microenvironment of maternal-fetal interface, but whether there are new clusters of macrophages with new characteristic need to be furtherly investigated.

Recurrent spontaneous abortion (RSA) is a common disease for women of childbearing age (Deroux, Dumestre-Perard, Dunand-Faure, Bouillet, & Hoffmann, 2017; Khalaj et al., 2016). At present, it is believed that in addition to fetal chromosomal abnormalities, increasing studies have found that the abnormal immune function of the mother’s decidua is also an important factor (Hu, Tang, Mor, & Liao, 2016; Kang, Zhang, & Zhao, 2016). At the maternal-fetal interface, the status of dMΦ inflammatory polarization changed with the development process of pregnancy, the balance of polarization between M1 and M2 macrophages is important for various processes of normal pregnancy, such as trophoblast invasion, spiral artery remodeling, and apoptotic cell phagocytosis (Buckley, Whitley, Dumitriu, & Cartwright, 2016; Li et al., 2016; Ning, Liu, & Lash, 2016),. Conversely, the dysregulated polarization of macrophages was associated with inadequate remodeling of the uterine vessels and defective trophoblast invasion and finally led to spontaneous abortion, preeclampsia and preterm birth (Cappelletti et al., 2017; Ferreira, Meissner, Tilburgs, & Strominger, 2017; Triggianese, Perricone, Chimenti, De Carolis, & Perricone, 2016),. So, it is very important to study the mechanism of the polarization of dMΦ during normal and pathological pregnancy.

Although increasing evidence has indicated the critical roles of dMΦ in pregnancy-related diseases (Liu et al., 2017), the properties of dysregulated macrophage polarization are still poorly understood. Here, we found that CD45^+^CD14^+^ dMΦ can divided into 13 cluster using t-SNE analysis, and normal pregnancy decidua have more percent composition of M1-type macrophages, but RSA patients have more percent composition of M2-type macrophages, and their expression profile are different from that of peripheral blood macrophages. We found that cluster 10 is the specialized cluster in normal pregnancy and cluster 7 highly expressed CXCL9/10/11, and identified many key transcription factors for each cluster. This will deepen our current knowledge of macrophage polarization involved in physiological or pathological pregnancy processes, and will allow us to develop therapies to improve pregnancy outcomes in the future.

## RESULTS

### scRNA-seq analysis of normal dMΦ

To elucidate the cells heterogeneity of dMΦ during normal pregnancy, we performed single-cell RNA-seq (scRNA-seq) on CD45^+^CD14^+^ dMΦ isolated by flow cytometry from decidual mononuclear cells (dMCs) in normal patient. We obtained a total of 14,701 single-cell transcriptomes (10,513 in Normal 1; 4,188 in Normal 2) after quality control (Supplmentary Figure 1 A-C). T-distributed stochastic neighbor embedding (t-SNE) visualization of the combined normal data revealed 17 major clusters based on their marker gene expression, which correspond to macrophages (PTPRC, CD14, MS4A7, CTSS), NKT (PTPRC, NKG7, NCAM1, CD3E, KLRD1), dendritic cells (DCs) (PTPRC, CD1C), Osteoclast & Macrophages (PTPRC, ACP5, RANK, CTSK), T cells (PTPRC, CD3E, CD8A), NK (PTPRC, NKG7, NCAM1, KLRD1), B & Plasma Cells (PTPRC, CD79A, MS4A1), smooth muscle cells (TAGLN、ACTA2、DCN、COL1A1) and endometrial epithelial cells (PAEP, GPX3 and SLPI) (Figures 1A-C and Supplmentary Table S1), and each cluster have its own specific marker genes (Figures 1D). We analyzed the percent composition of all kinds of cell types in both Normal 1 and Normal 2 (Supplmentary Figure 1D). Macrophages occupied about 82.7% of the total cells in the individual sample (Supplmentary Figure 1E), and we next focused on analyzing the function and characterization of dMΦ.

**Figure 1.**
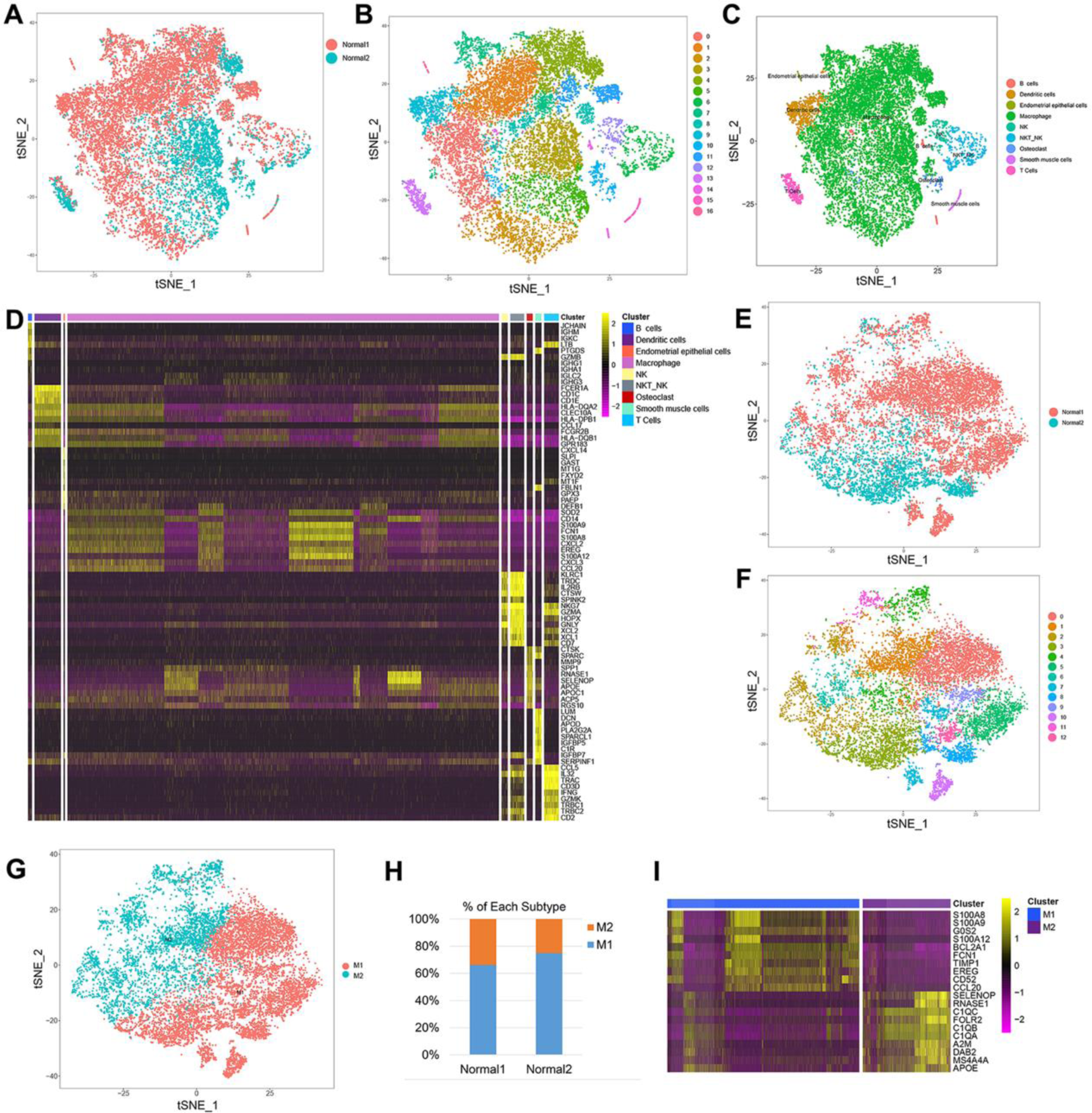
A single-cell analysis of normal human decidual mononuclear cells. (A) t-Distributed stochastic neighbour embedding (t-SNE) analysis of decidual mononuclear cells from the combined Normal 1 and Normal 2 sample. (B) Seventeen healthy human decidual cell clusters. t-SNE analysis of 14,701 decidual mononuclear cells and colored by clustering. (C) Cell type analysis of decidual mononuclear cells annotated by marker genes and colored by cell types. (D) Heatmap analysis of the expression of specific marker genes for each cluster. (E) t-SNE analysis of decidual macrophage from the combined Normal 1 and Normal 2 sample. (F) Thirteen healthy human decidual macrophage clusters. t-SNE analysis of 12,470 decidual macrophage and colored by clusters. (G) M1/2 type analysis of decidual macrophage annotated by marker genes and colored by cell types. (H) Percent composition analysis of M1/2 cells occupied in Normal 1 and Normal 2. (I) Heatmap analysis of the expression of M1/2 marker genes.

Macrophage cell types were furtherly analyzed in-depth by identifying sub-clusters. After re-clustering, 12,470 macrophage cells (9,005 in Normal 1; 3,465 in Normal 2) in 13 clusters were detected in normal decidua samples (Figure 1E, F and Supplmentary Figure 1F-H). Due to that macrophages can be polarized into classically activated (M1) and alternatively activated (M2) macrophages, based on the polarization markers of dMΦ from previously reported (Vento-Tormo et al., 2018) (M1 marker: S100A9, CD68, CD163, CSF1R, IL1β, TNF, IL6, CXCL2, CXCL3, CXCL8; M2 marker: IL10, MRC1, MAF, F13A1, C3, C1QA, C1QB, C1QC, IGF1, IGFBP4), we divided our dMΦ into two parts, M1 cells (5961 in Normal 1 and 2596 in Normal 2) and M2 cells (3044 cells in Normal 1 and 869 cells in Normal 2) (Figure 1G and Supplmentary Table S2). Interestingly, the percentage composition of M1 cells (70.6%) were significantly higher than M2 cells (29.4%) in normal decidua (Figure 1H), which is different from the traditional opinion that the higher composition of M1 cells is harmful to normal pregnancy. We also found that the specific marker gene of M1 and M2 (Figure 1I). These observations strongly suggest that normal pregnancy is linked to the relatively high polarization level of M1 macrophages.

For further characterization of M1 or M2 cells in normal decidua, we used Database for Annotation, Visualization and Integrated Discovery (DAVID) to identify KEGG and GO terms enriched among marker genes of M1 or M2. We observed that M1 macrophages might take part in a set of ribosome translational biological processes and several cellular signaling pathways (Ribosome, NOD-like receptor, NF-kappa B and TNF signaling pathway) (Supplementary Figure 2A-C and G), whereas M2 macrophages might take part in a set of antigen processing and presentation biological processes and several cellular signaling pathways (Antigen processing and presentation, lysosome and phagosome signaling pathway) (Supplementary Fig. 2D-F and H). Thus, these results suggest that the marker genes in M1 or M2 play different biological functions in the decidua of early pregnancy.

### scRNA-seq analysis of normal and RSA dMΦ

To elucidate the dynamic changes of dMΦ during RSA pathogenesis, we performed scRNA-seq on CD14^+^CD45^+^ dMΦ isolated from RSA patients. We obtained a total of 13,879 single-cell transcriptomes (9,889 in RSA1, 3990 in RSA2) after removing low quality cells (Supplmentary Figure 3A-C). After combined with the Normal 14,701 single-cell transcriptomes, T-distributed stochastic neighbor embedding (t-SNE) visualization of the combined normal and RSA data also revealed 17 major clusters, which are consistent with the Normal cells cluster based on marker gene expression (Figures 2A-C and Supplementary Table S3), and each cluster has their specific marker genes (Figure 2D). We analyze the percent composition of all kinds of cells in the four samples (Supplementary Figure 2D). Among these cell types, macrophages occupied 80.3% of the total cells in the individual sample (Supplementary Figure 2E), and we next focused on analyzing the function and characterization of macrophages.

**Figure 2.**
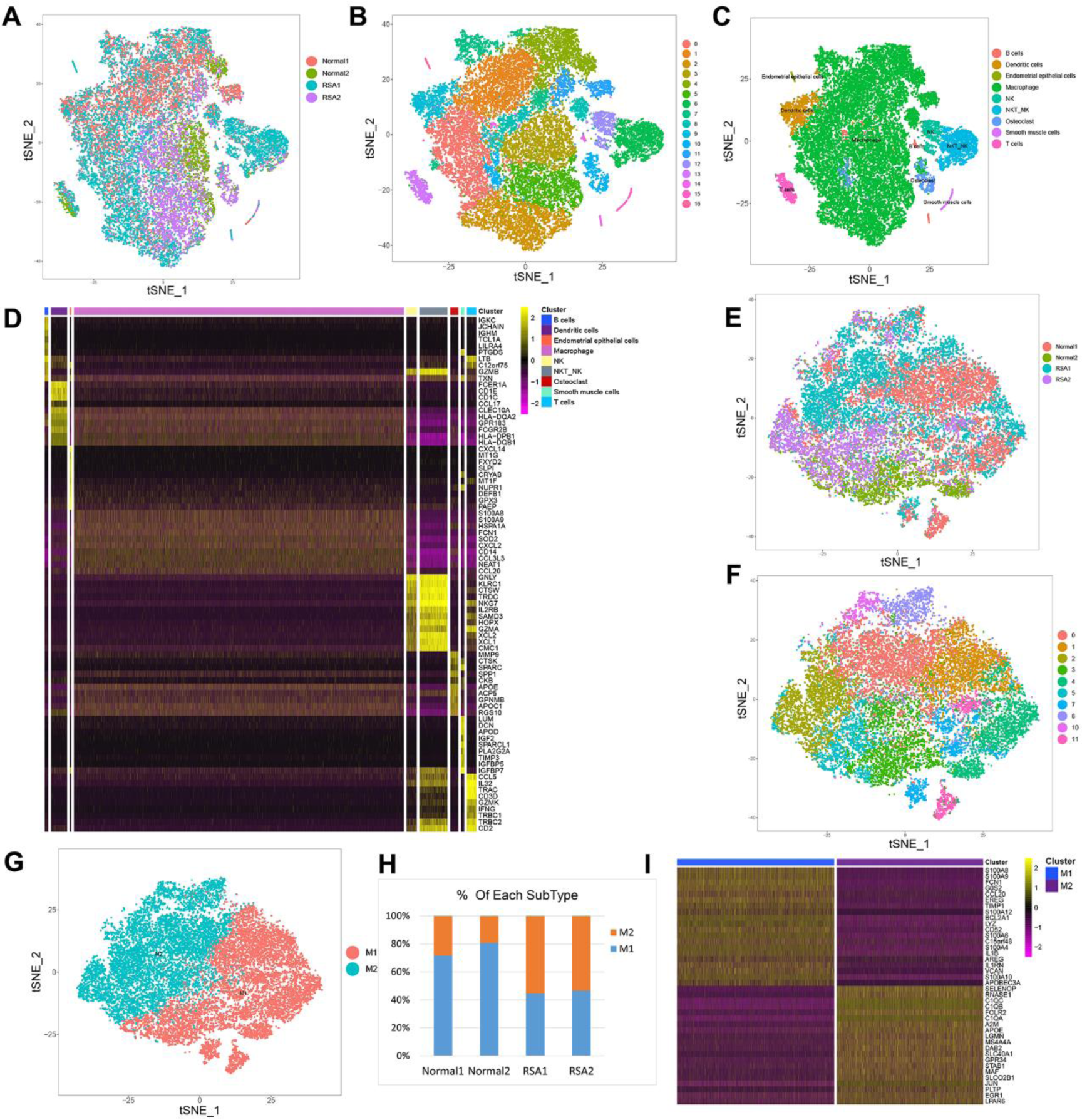
A single-cell analysis of combined normal and RSA human decidual mononuclear cells. (A) t-SNE analysis of decidual mononuclear cells from the combined four samples (Normal 1, Normal 2 and RSA1, RSA2). (B) Seventeen human decidual cell clusters. t-SNE analysis of 28,580 decidual mononuclear cells from (A) and colored by clusters. (C) Cell type analysis of decidual mononuclear cells annotated by marker genes and colored by cell types. (D) Heatmap analysis of the expression of specific marker genes for each cluster. (E) t-SNE analysis of decidual macrophages from the combined four samples (Normal 1, Normal 2 and RSA1, RSA2). (F) Thirteen human decidual macrophage clusters. t-SNE analysis of 23,062 decidual macrophages and colored by clustering. (G) M1/2 type analysis of decidual macrophages annotated by marker genes and colored by cell types. (H) Percent composition analysis of M1/2 cells occupied in Normal 1, Normal 2 and RSA1, RSA2. (I) Heatmap analysis of the expression of M1/2 marker genes in the combined four samples.

**Figure 3.**
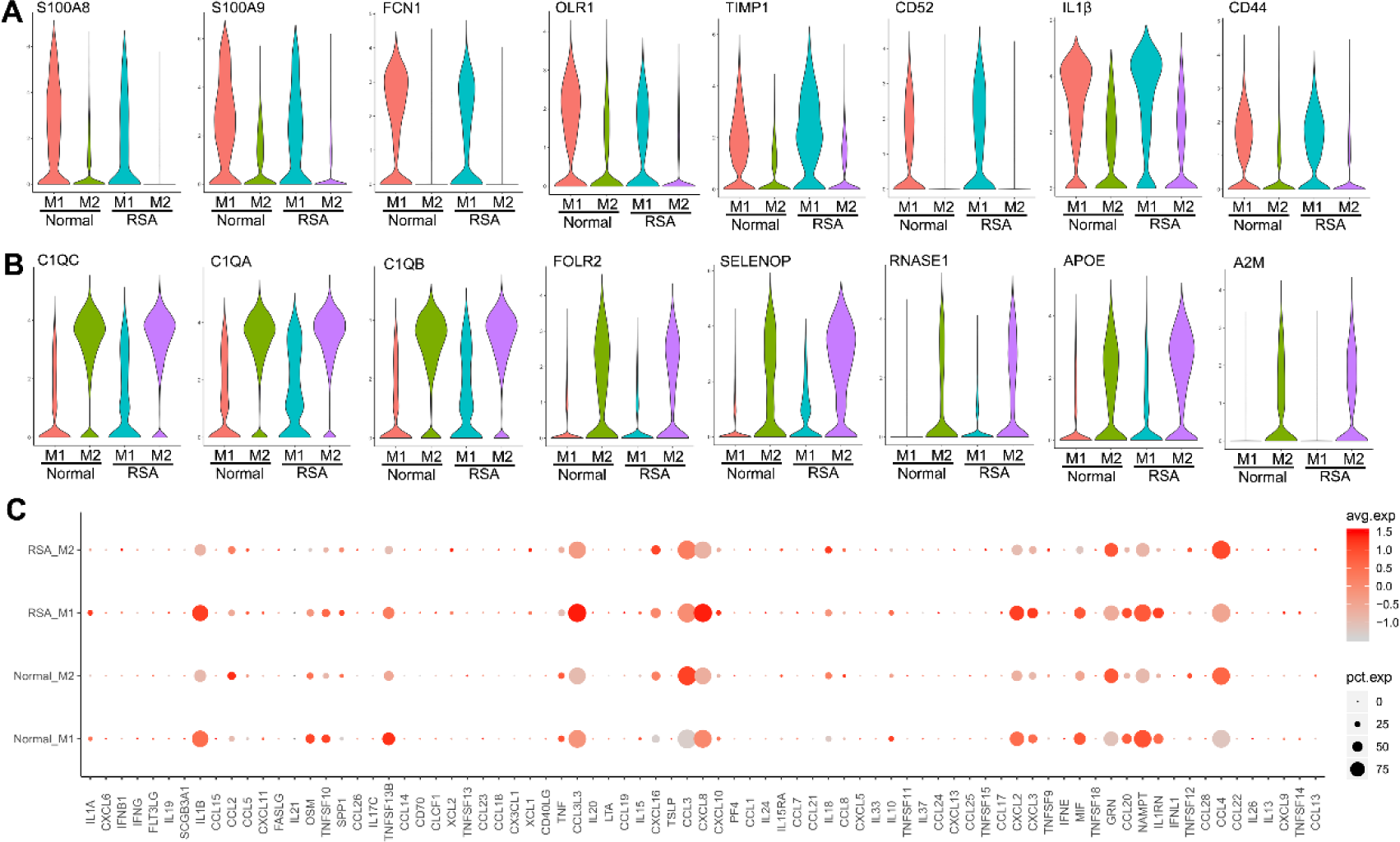
Identification of the new markers and secreted cytokines of the dMΦ. (A, B) Violin plot showing the new markers of M1 macrophages (A) and M2 macrophages (B). (C) Gene bubble plot showing the secreted cytokines and chemokines of M1 or M2 macrophages.

Macrophage cell types were furtherly analyzed in-depth by identifying sub-clusters. After re-clustering, multiple smaller new clusters emerged, 23,062 macrophage cells (9,005 in Normal 1; 3,465 in Normal 2; 6,993 in RSA1; 3,599 in RSA2) in 13 clusters were detected in normal and RSA decidua samples (Figure 2E, F and Supplementary Figure 3F-H). Based on the macrophage polarization markers from decidua previously reported (Vento-Tormo et al., 2018), we also divided our dMΦ into two parts, M1 cells (5961 in Normal 1, 2596 in Normal 2, 2271 in RSA 1, 1113 in RSA 2) and M2 cells (3044 cells in Normal 1, 869 cells in Normal 2, 4722 RSA 1, 2486 RSA 2) (Figure 2G and Supplementary Table S4). Surprisingly, we observed that the percentage composition of M1 cells in the RSA decidua was significantly lower than that of the normal decidua (31.7% *vs*. 70.6%), while the percentage composition of M2 cells in the RSA decidua were significantly higher than that of the normal decidua (68.3% *vs.* 29.4%) (Figure 2H). We also found the specific marker genes of M1 and M2-type macrophages (Figure 2I). These results are contrary to that composition of normal pregnancy, and strongly indicate that M1 polarization level of macrophages plays a very important role in maintaining the normal pregnancy, and much more M2 polarization of macrophages maybe the reason of the RSA pregnancy.

For further characterization of M1 or M2 cells in RSA decidua, we used DAVID to identify KEGG and GO terms enriched among marker genes of M1 or M2. We observed that M1 macrophage might take part in a set of inflammatory response and immune system process biological processes and several cellular signaling pathways (TNF and NOD-like receptor signaling pathway) (Supplementary Figure 4A-C and G), whereas M2 macrophage might take part in a set of immune response biological processes and several cellular signaling pathways (antigen processing and presentation, staphylococcus aureus infection and lysosome signaling pathway) (Supplementary Figure 4D-F and H). Thus, these results suggest that the characterization of M1 or M2 macrophage from normal or RSA is different, and further reflect the different biological function of M1 and M2 in the decidua of early pregnancy.

Next, we analyzed genes that were differentially expressed in macrophages of normal and RSA decidua. Overall, we observed that numbers of genes were upregulated (102 genes) and downregulated (122 genes) in the RSA group compared with the normal group (Supplementary Table S5). To gain insights into the potential functional differences of the dynamic genes between the two groups, we performed functional enrichment analysis of the genes that were up- or down-regulated in normal and RSA groups. The upregulated genes (Top 5: TMSB4X, RPL18A, RPL39, S100A11, SH3BGRL3) (Supplementary Figure 4I) in the normal group were enriched for the functional categories of Ribosome, Phagosome and HIF-1 signaling pathway, indicating the activation of hypoxia induced genes in early pregnancy. The upregulated genes (Top 5: SELENOP, RNASE1, HLA-DQA1, GNLY, PDK4) (Supplementary Figure 4J) in the RSA group were enriched for the functional categories of Complement and coagulation, Toll-like receptor signaling and Estrogen signaling pathway, indicating the inhibition of inflammatory genes in macrophage of early pregnancy, whether this is related to the RSA requires furtherly study.

### New markers and secreted cytokines of the dMΦ

Previous studies labeled dMΦ with markers of peripheral blood macrophages, including classically activated (M1) markers (such as, TLR-2, TLR-4, CD80, CD86, iNOS, and MHC-II (HLA-DRB1)) and alternatively activated (M2) markers (mannitol receptor (ITGAM), CD206 (MRC1), CD163, CD209) (Brown, von Chamier, Allam, & Reyes, 2014). We detected the expression of these molecular in our single cell database, and found that the expression of these molecular (CD80, CD86, ITGAM, CD206 (MRC1), CD163 and CD209) are very low in the dMΦ and they cannot present the polarization state of decidua M1 or M2 (Supplementary Figure 5 A, B), which indicated that dMΦ isolated with classical markers of peripheral blood macrophages cannot reflect the real state of dMΦ in the maternal-fetal microenvironment.

Is there a more accurate marker for dMΦ in maternal-fetal interface microenvironment? We next intend to examine the new marker of dMΦ, and found that new markers (PTPRC, CD14, MS4A7 and CTSS) can represent the dMΦ (Supplementary Table S1). We also identified the M1 marker genes (S100A8, S100A9, G0S2, S100A12, BCL2A1, FCN1, TIMP1, EREG, CD52 and CCL20) and M2 marker genes (SELENOP, RNASE1, C1QC, FOLR2, C1QB, C1QA, A2M, DAB2, MS4A4A and APOE) of dMΦ, which are easier to distinguish the polarization states of macrophages (Figure 3A, B). These results indicated that the genes (PTPRC, CD14, MS4A7 and CTSS) maybe the ideal new potential markers to isolate the macrophage from the decidua tissue, and M1 or M2 marker genes are easier than the traditional markers to distinguish the phenotypes of macrophages.

We have known that classically M1 and M2 macrophage cells can release various cytokines and chemokines (such as M1: TNFα, IL1α, IL1β, IL6, IL12, CXCL9 and CXCL10; M2: IL10, TGFβ, CCL1, CCL17, CCL18, CCL22 and CCL24) respectively (Brown et al., 2014). However, we find that all these cytokines are not highly expressed in dMΦ, and their expression have no significantly difference between M1 and M2, except for IL-1β, and TGF-β seem more likely highly expressed in M1, but not M2 (Supplementary Figure 5C, D). We found that M2 type dMΦ cells highly expressed CCL3, CCL4, GRN, and M1 type cells highly expressed IL1β, TNFSF13B, CCL3L3, CXCL8, CXCL2, MIF, CCL20, IL1RN and NAMPT (Figure 3A-C). These results indicated that cytokines and chemokines released by dMΦ are different from those of peripheral blood macrophages.

### Cluster 10 as the specific cluster of dMΦ in normal group

We believed that dMΦ dysfunction is one of the important reasons that caused RSA, so we speculated that there maybe have some specific clusters disappeared in RSA group compared to normal group. After merged analysis of RSA and normal group by t-SNE, we found that cluster 10 almost disappeared in RSA group (Figure 2E), and this cluster mainly expressed marker genes (ZFAND2A, BAG3, HSPD1 and CACYBP) genes (Figure 4A) and highly expressed cytokines and chemokines (IL1β, CCL3L3, CCL3, CXCL 8, CXCL2, MIF, NAMPT, IL1RN and CCL4), which specifically expressed TNFSF14 (Figure 4B). To gain insights into the potential functional of this cluster, we performed DAVID software to identify KEGG and GO terms enriched among marker genes of cluster 10. We observed that cluster 10 might take part in a set of cellular component movement and response to unfolded protein process biological processes (Supplementary Figure 6A) and several cellular signaling pathways (glycolysis/gluconeogenesis, ribosome and HIF-1 signaling pathway) (Supplementary Figure 6B). Thus, these results suggest that cluster 10 may take part in regulating hypoxia response or other stimuli, and may play an important role in the implantation process of embryo.

**Figure 4.**
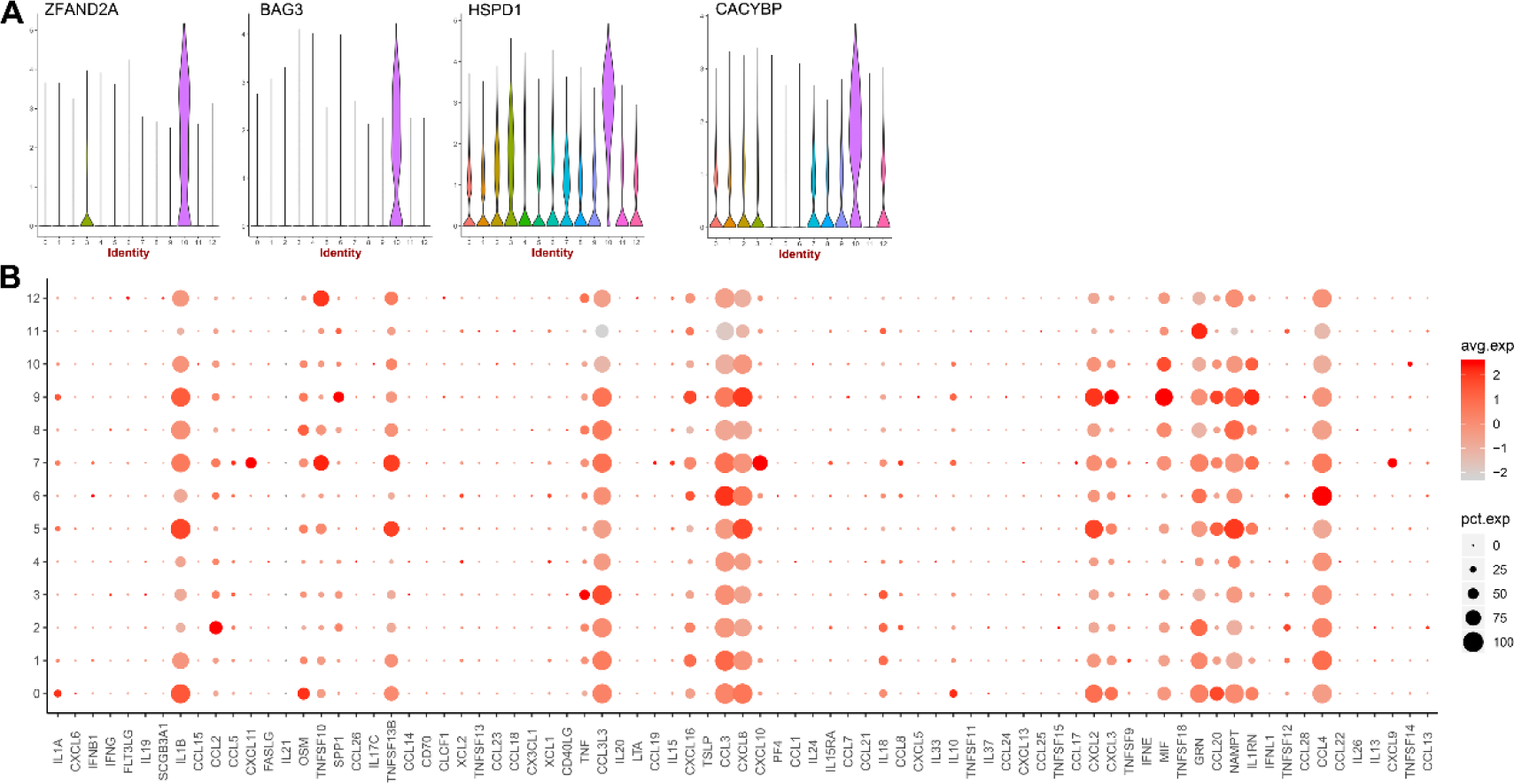
Characterization of Cluster 10 as the specific cluster of dMΦ in normal group. (A) Violin plot showing the marker genes of cluster 10. (B) Gene bubble plot showing the secreted cytokines and chemokines of the clusters.

### Cluster 7 specifically highly expressed CXCL9/10/11 chemokines

Next, we speculated that there might be some specific clusters in decidua tissue which secrete specific cytokines to maintain pregnancy. After merged analysis of RSA and normal group by t-SNE, we found that although cluster 7 displayed with no difference in cell number between Normal and RSA group (Supplementary Table S4), but this cluster specifically expressed chemokines of CXCL9/10/11 (Figure 5A) and mainly expressed marker genes (GBP1, ISG20, TNFSF10, IFIT3, IFITM1, ISG15, IDO1 and IL4I1) (Figure 5B). To gain insights into the potential functional of this cluster, we performed DAVID software to identify KEGG and GO terms enriched among marker genes of cluster 7. We observed that cluster 7 might take part in a set of biological processes and several cellular signaling pathways, such as type I interferon signaling pathway, cytokine-mediated signaling pathway and immune system process (Supplementary Figure 7A, B). Thus, these results suggest that cluster 7 mainly take part in regulating antiviral immune response.

**Figure 5.**
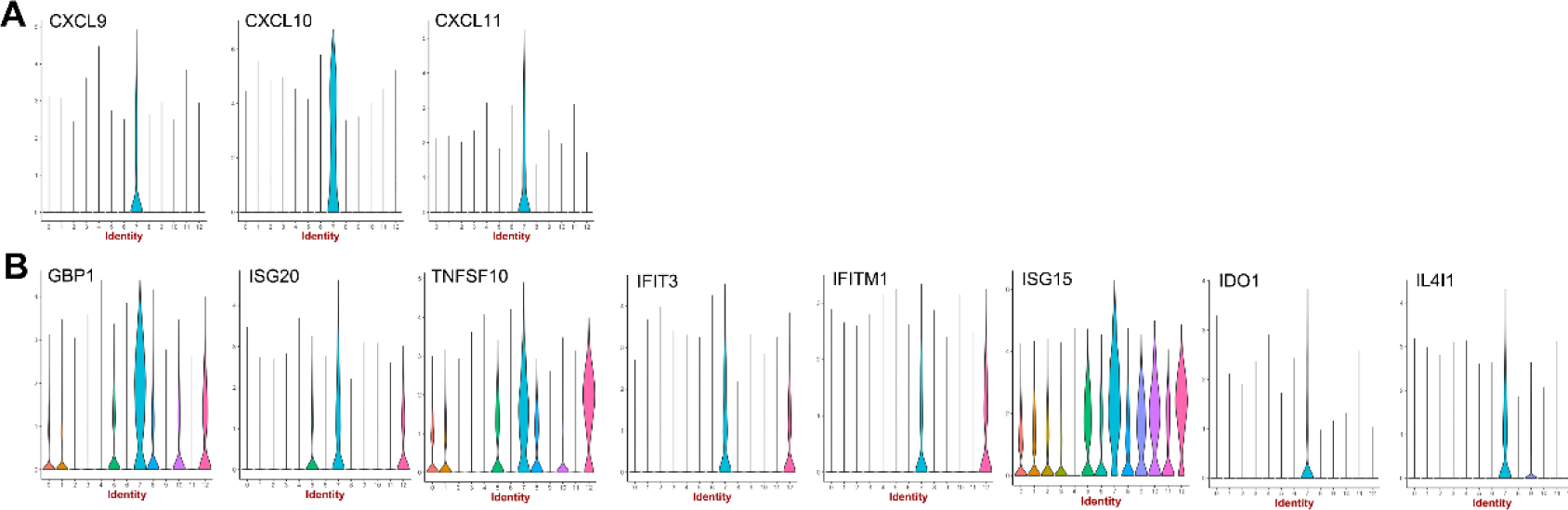
Characterization of Cluster 7 dMΦ in normal group. (A, B) Violin plot showing the secreted chemokines and the marker genes of cluster 10.

### Single-cell trajectory analysis of dMΦ

We then ordered individual macrophage cells by Monocle2 to pseudo-time trajectory analysis to reconstruct their differentiation relationship. We found that the macrophage from normal and RSA decidua formed a continuous “V-shaped” trajectory with 7 states (Figure 6A and Supplementary Figure 8A, B). Among them, states 1, 2 and 3 were occupied by M1 type macrophages which included cluster 0, 5, 7, 8, 9, 10, 12 and state 5 was occupied by M2 type macrophage which included cluster 1, 2, 3, 4, 6, 11. M1/M2 macrophage mainly occupied the two heads and mixed macrophages at the turn and tail (Figure 6B and Supplementary Figure 8C, D). States 1, 2 and 3 take part in Ribosome, Parkinson’s disease and inflammation related signaling pathway, while state 5 takes part in antigen processing and presentation signaling pathway (Supplementary Figure 8E-I). The M1/M2 macrophage branch further bifurcated into two subbranches (Figure 6A). One subbranch (State 6) was occupied by cells with highly expressed MALAT1, NEAT1 and S100A6 genes which involved in antigen processing and presentation signaling pathway (Supplementary Figure 8J), whereas the other (State 7) showed strong expression of genes (e.g., HLA-DPB1, HLA-DRA, CLEC10A and RPL15) involved in ribosome signaling pathway (Supplementary Figure 8K). We also found that development process of dMΦ was from M1 to M2 in the early pregnancy (Figure 6C). In short, our dataset enabled the delineation of the macrophage lineage relationship and shed light on the regulatory mechanism of lineage development. These expression patterns were consistent with two different fates of the trajectory, indicating that our scRNA-seq analysis correlated with the macrophage developmental process.

**Figure 6.**
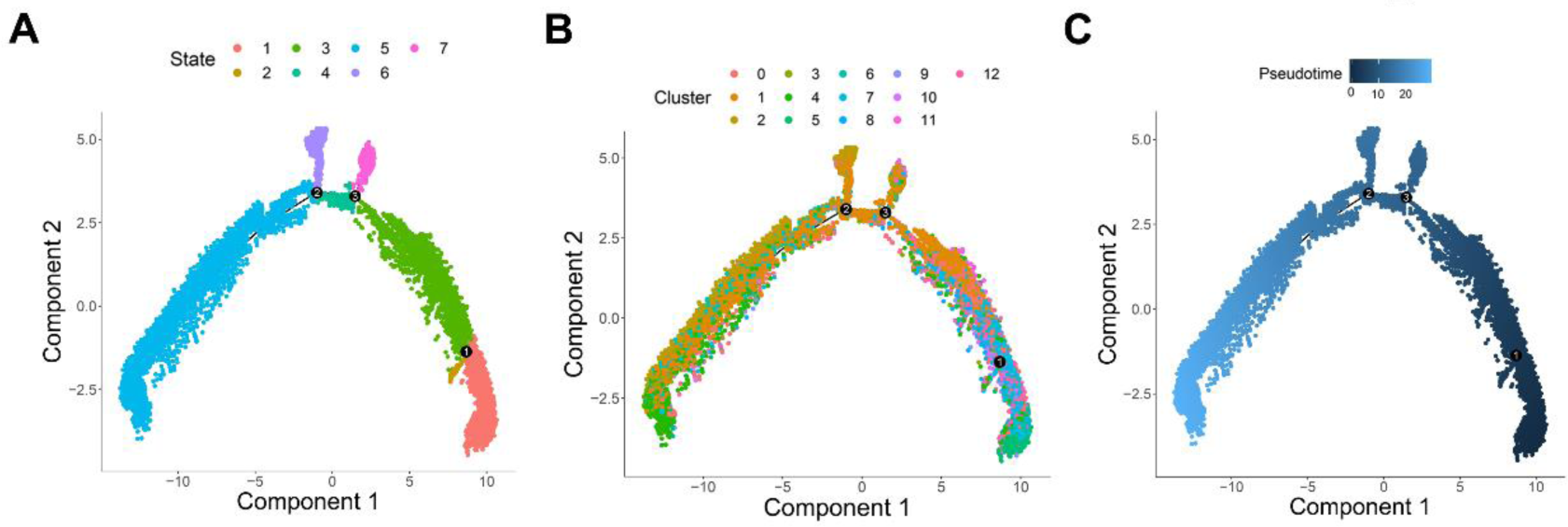
Single-cell trajectory analysis of decidual macrophages. (A-C) Monocle pseudotime trajectory showing the progression of dMΦ.

### Key transcription factors of dMΦ clusters

Transcription factors (TFs) play key regulatory roles in many biological processes. It is reported that NFκB, STAT1, STAT5, IRF3, and IRF5 as the M1 key transcription factors, and STAT6, IRF4, PPARδ, and PPARγ as the M2 key transcription factors, which have been shown to regulate the expression of M1 or M2 genes. But in our present study, we found that these transcription factors expressed lowly in dMΦ and had no specificity to M1 or M2 type, except for NFκB and STAT1 TFs. It seems that NFκB may be the major TFs involved in M1 macrophage polarization, and other classic transcription factors maybe have little effect on macrophage polarization (Supplementary Figure 9A-H).

To explore central factors that might contribute to regulate the specific gene expression of each cluster, we firstly investigated possible upstream TFs for these specific genes using TF-binding motif enrichment analysis. We found that top 10 TFs can regulate the gene transcription of normal or combined normal and RSA macrophage (Figure 7A, B). We identified CEBPB, FOSL2, FOXO3, IRF7, NFκB1, PRDM1, MYC and MAFF as the key transcriptional factors for M1 type macrophage including cluster 0, 5, 7, 8, 9, and 10, while EGR1 and TCF12 as the key transcriptional factors for M2 type macrophage, including cluster 2, 6 and 11 (Figure 7C). Therefore, these findings suggested that TFs might play a central role in regulating expression of during human preimplantation development.

**Figure 7.**
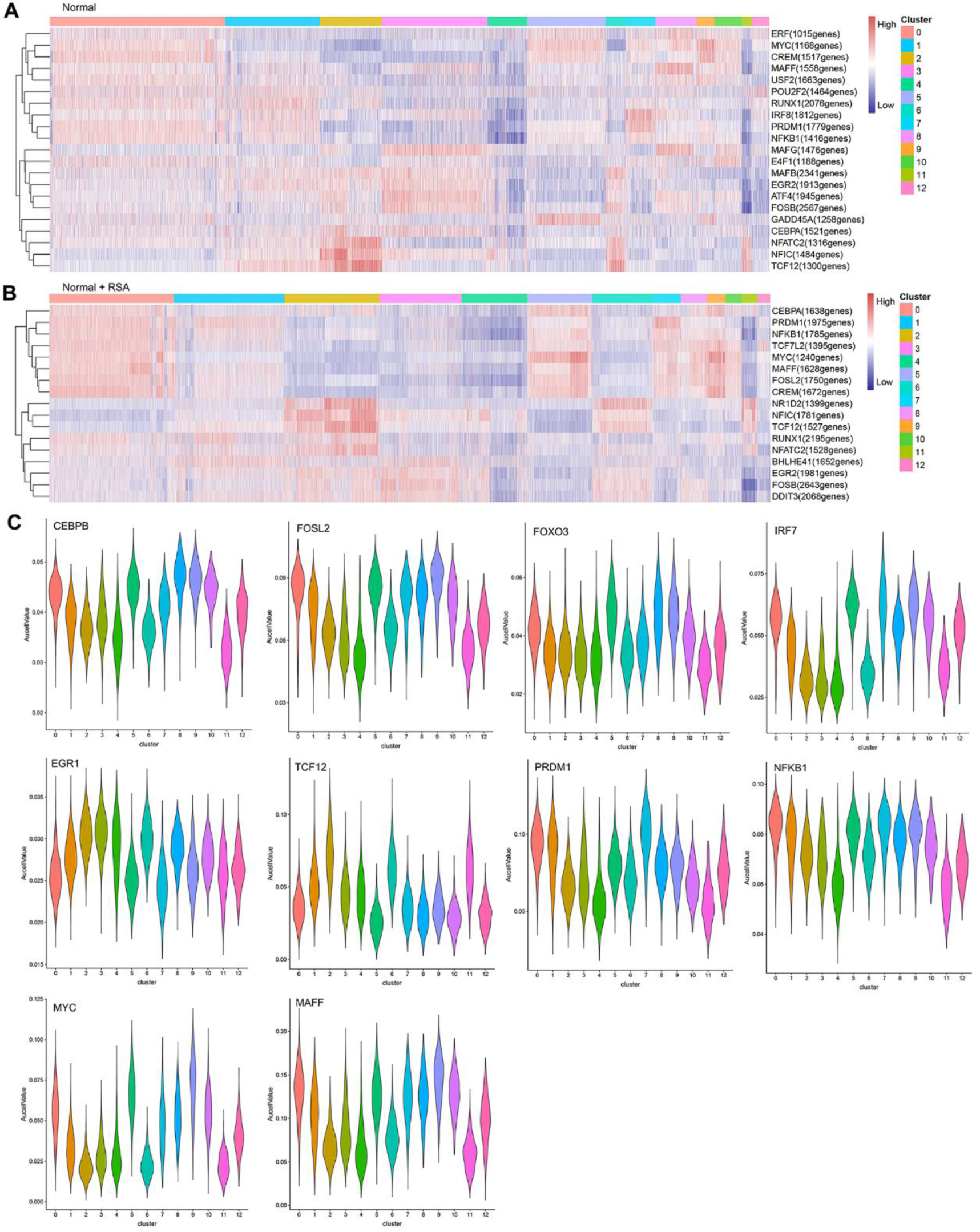
Identification of the key transcription factors of dMΦ. (A, B) Heatmap analysis of the expression of transcription factor in normal dMΦ (A) and the combined four samples (B). (C) Violin plot showing the expression of transcription factors in each cluster. Supplementary material for “Single cell transcriptome analysis of decidua macrophage in normal and recurrent spontaneous abortion patient”

## DISCUSSION

In this report, we found that normal pregnancy decidua had more percent composition of M1 macrophage, and RSA patient decidua had more percent composition of M2 macrophage, and their characteristics (such as, mRNA expression profile, secreted cytokines and chemokines or transcription factors) were different from those of peripheral blood macrophages. These results will deepen our current knowledge of macrophage activation involved in physiological or pathological pregnancy processes.

The marker genes for isolating dMΦ are very important for studying dMΦ functions. In many previous studies, the surface marker genes of macrophages from peripheral blood, which characterized by the expression of the mannose receptors CD206, CD80, CD209, CD16 or CD163, were utilized as FACS marker to isolate dMΦ, but dMΦ had tissue specificity, and these markers did not separate well for dMΦ (Bockle, Solder, Kind, Romani, & Sepp, 2008; Kammerer et al., 2003; Ono et al., 2018). For exploring more precise macrophage types, we isolated the dMCs using CD45^+^CD14^+^ antibodies, and we found 17 clusters with 9 types of cells. dMΦ occupied up to about 82%. We found that CD45^+^CD14^+^MS4A7^+^ might be the better combined markers for isolating dMΦ. Furthermore, we identified the new marker genes for dMΦ, but these specific markers need to be further identified.

During early pregnancy, the polarization of dMΦ for success pregnancy is controversial (Ding et al., 2020; Jena, Nayak, Chen, & Nayak, 2019). It is reported that the polarization of dMΦ to an anti-inflammatory state is critical for the success of the pregnancy (Ben Amara et al., 2013). Decidual macrophage activation towards a more M1 phenotype has been associated with the pathology of RSA (Tsao, Wu, Chang, Wu, & Ho, 2018). But we found that that the percentage composition of M1 cells in the RSA decidua were significantly lower than M1 cells in the normal decidua (31.7% *vs.* 70.6%), while the percentage composition of M2 cells in the RSA decidua were significantly higher than M2 cells in the normal decidua (68.3% *vs.* 29.4%). These results indicate that polarization level of M1 macrophage play a very important role in maintaining the normal pregnancy, and much more polarization of M2 macrophage may be harmful for the early pregnancy. However, our understanding of the function and polarization of dMΦ is still limited and needs to be studied in the future.

The non-pregnant endometrium has a low concentration of macrophages, with fluctuating numbers along the menstrual cycle (Huhn et al., 2020). After fertilization, macrophages represent the second most abundant decidual leukocyte population (around 20–30%) (Yang, Zheng, & Jin, 2019). Human dMΦ may be involved in remodeling the endometrium, trophoblast invasion, and the development of the tolerant milieu required for successful progression of pregnancy (Garcia-Gomez et al., 2019). Macrophage can respond efficiently to environmental signals and change their phenotype and physiology in response to cytokines and extra stimulation (Olmos-Ortiz et al., 2019). According to some reports, dMΦ represent alternatively activated M2 macrophage, particularly since local decidual microenvironments are favorable for alternative activation (Ticconi, Pietropolli, Di Simone, Piccione, & Fazleabas, 2019). The M1 phenotype typically expressed IL-1β, TNF-a, IL-12 and lowly expressed IL-10, whereas the M2 macrophage phenotype typically expressed IL-10 and TGF-β, but lowly expressed IL-12 (Liu et al., 2017; Zhang, He, Wang, & Liao, 2017). But in our study, we found that M2 type cells highly expressed CCL3, CCL4, GRN, and M1 type cell highly expressed IL1β, TNFSF13B, and CCL3L3. Previously, it was reported that an inflammatory M1 phenotype was associated with human implantation, characterized by the upregulation of TNF-α, MIP-1β and GRO-α, together with the secretion of remodeling factors such as matrix metalloproteinases (MMP) -3 and -9, suggesting that decidual M1 macrophages modulate an inflammatory environment that results in increased trophoblastic invasion and spiral artery remodeling.

We found that Cluster 10 as the specific cluster of dMΦ in normal group and almost disappeared in RSA group, and found that this cluster highly expressed inflammation cytokines and chemokines. Cluster 10 might take part in HIF-1 signaling pathway, which suggests that cluster 10 may take part in regulating hypoxia response or other stimuli, and may play an important role in the implantation process of embryo (Macklin, McAuliffe, Pugh, & Yamamoto, 2017; Soares, Iqbal, & Kozai, 2017), We also found that cluster 7 specifically expressed chemokines of CXCL9/10/11 and mainly expressed marker genes of type I interferon signaling pathway, which suggests that cluster 7 mainly take part in regulating antiviral immune response. Whether these clusters are related to the miscarriages requires further study.

## MATERIAL AND METHODS

### Isolating mononuclear cells from decidua

Decidua samples were collected from RSA patients or control women undergoing elective terminations of first trimester pregnancies. To isolate decidual mononuclear cells (dMCs), decidual tissue pieces were first washed, remaining tissue was then minced and digested by Collagenase IV (0.1 g/100 mL, sigma) and DNase I (150 U/mL, Sigma) for 40 min at 37°C whilst being gently shaken. The collagenase was quenched by addition of DMEM/F12 with 10% FCS and then filtered through 100 μm and then 40 μm filters. dMCs were isolated following a percoll gradient centrifugation step. This protocol has been adapted from previous reports (Vishnyakova, Elchaninov, Fatkhudinov, & Sukhikh, 2019).

### Cell staining and data acquisition by flow cytometry

dMCs were washed in complete medium. Cells were then counted and distributed at approximately 1×10^6^ cells/mL in 200 μL FACS wash (FW, 1×PBS, 2% FCS). Cells were stained with CD14 and CD45 antibody for 15 min at 4 °C. Viability was assessed using DAPI staining incubated for 5 min at 4 °C. Cells were then analyzed on a LSR Fortessa (BD Biosciences). Macrophages were gated on as live CD14^+^CD45^+^ lymphocytes. Further gating on subsets was done as indicated in figure legends. The following antibodies and reagents were used: DAPI (Life Technologies), FITC-CD14 (Biolegend), Cy5-CD45 (BioLegend).

### Single-Cell RNA Sequencing (scRNA-seq)

Cells were prepared according to the 10×Genomics® Cell Preparation Guide. The protoplast suspension was loaded into Chromium microfluidic chips with chemistry and barcoded with a 10×Chromium Controller (10×Genomics). RNA from the barcoded cells was subsequently reverse-transcribed and sequencing libraries constructed with reagents from a Chromium Single Cell 30 reagent kit (10×Genomics). Sequencing was performed with Illumina (HiSeq 2000) according to the manufacturer’s instructions. All single cell sequencing database deposited under GEO Accession number (GSE169269).

### Single-cell RNA statistical analysis

We applied fastp with default parameters to filter the adaptor sequence and remove the low-quality reads to obtain clean data (Rosenberg et al., 2018). UMI-tools were applied for single cell transcriptome analysis to identify the cell barcode whitelist (Han et al., 2018). The UMI-based clean data was mapped to the human genome (Ensemble version 91) utilizing STAR (Picelli et al., 2014) mapping with customized parameters from the UMI-tools standard pipeline to determine the UMIs counts of each sample. In order to minimize the sample batch, we applied down sample analysis to the samples sequenced according to the mean reads per cell of each sample and achieved a cell expression table with a sample barcode. Cells containing over 200 expressed genes and a mitochondria UMI rate below 20% passed the cell quality filtering and mitochondria genes were removed in the expression table but used for cell expression regression to avoid the effect of the cell status for clustering analysis and marker analysis of each cluster. Seurat package version 2.3.4 (https://satijalab.org/seurat/) was used for cell normalization and regression based on the expression table according to the UMI counts of each sample and the percentage mitochondria rate to obtain the scaled data. PCA was constructed based on the scaled data with all highly variable genes. The top 8 principals were used for tSNE construction. Utilizing the cluster method (normal group: k-mean method and K=7; normal group and RSA group: graph cluster method and resolution=1), we acquired the cell cluster result based on the PCA of the top 8 principals and calculated the marker genes using the Find All Markers function with the Wilcoxon Rank-Sum Test under following criteria: LogFC>0.25; p<0.05; min.pct>0.1.

### Pseudotime analysis

We applied Single-Cell Trajectories analysis using Monocle2 (http://cole-trapnelllab.github.io/monocle-release) with DDR-Tree and default parameters. Before Monocle analysis, we selected the marker genes of the Seurat clustering result. The raw expression counts of the cell passed filtering. Based on the pseudo-time analysis, branch expression analysis modelling (BEAM Analysis) was applied for branch fate determined gene analysis (Vishnyakova et al., 2019).

### SCENIC analysis

The SCENIC analysis was run on the cells that passed the filtering, using the 20 thousand motifs database for RcisTarget and GRNboost, as previously described (Sun et al., 2020).

### Pathway analysis

Pathway analysis was used to identify the significant pathway of the differential genes according to the KEGG database(Zheng, Hou, Zhou, Li, & Cao, 2017). We used Fisher’s exact test to select the significant pathway, where the threshold of significance was defined by the P-value and FDR. The cases were selected when p<0.05.

## ACKNOWLEDGEMENTS

This study was supported by the Key Program of the National Natural Science Foundation of China (81730039), the National Natural Science Foundation of China (82071653, 81671460, 81871167, 82071644, 81971384), the Natural Science Foundation of Shanghai (18ZR1430000), the National Key Research and Development Program of China (2017YFC1001401), and Shanghai Municipal Medical and Health Discipline Construction Projects (2017ZZ02015) to Liping Jin.

## AUTHOR CONTRIBUTIONS

Liping Jin and Xiaoping Wan designed the experiments. Fenglian Yang and Yongbo Zhao collected the patient samples with assistance from Haili Gan. Qingliang Zheng and Xiang-Hong Xu carried out the experiments and analyzed the data and interpreted the findings. Liping Jin and Qingliang Zheng wrote the manuscript.

---

**Figure S1.**
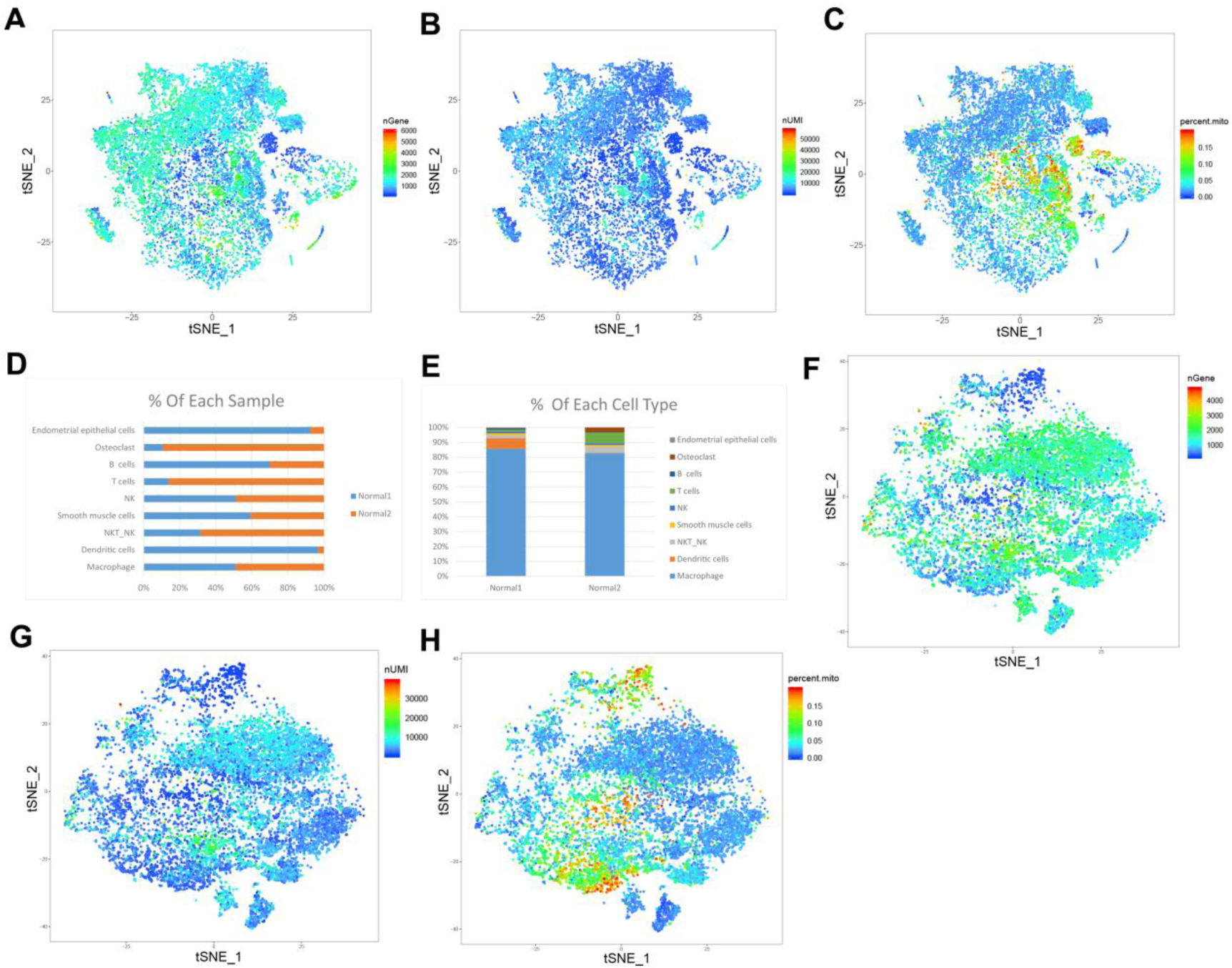
A single-cell analysis of normal human decidual mononuclear cells. (A-C) t-Distributed stochastic neighbour embedding (t-SNE) analysis of decidual mononuclear cells from the combined Normal 1 and Normal 2 sample on nGene (A), nUMI (B) and pecernt mitochondrion (C). (D, E) Percent composition analysis of each cell type in Normal 1 and Normal 2 sample. (F-H) t-SNE analysis of decidual macrophage cells from the combined Normal 1 and Normal 2 sample on nGene (F), nUMI (G) and pecernt mitochondrion (H).

**Figure S2.**
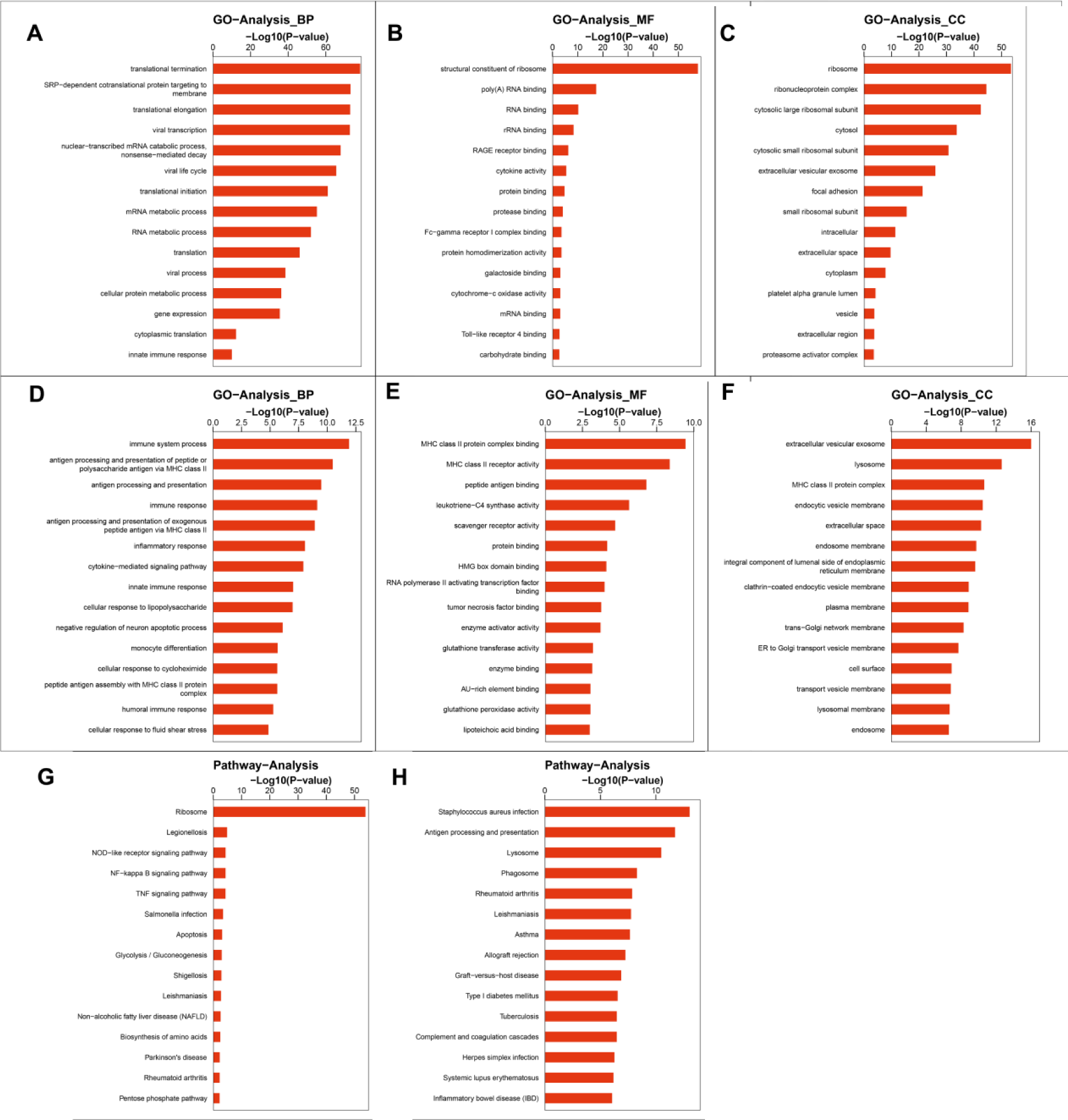
(A-F) GO analysis of the marker genes in M1 (A-C) or M2 (D-F) macrophage. (G,H) KEGG analysis of the marker genes in M1 (G) or M2 (H) macrophage.

**Figure S3.**
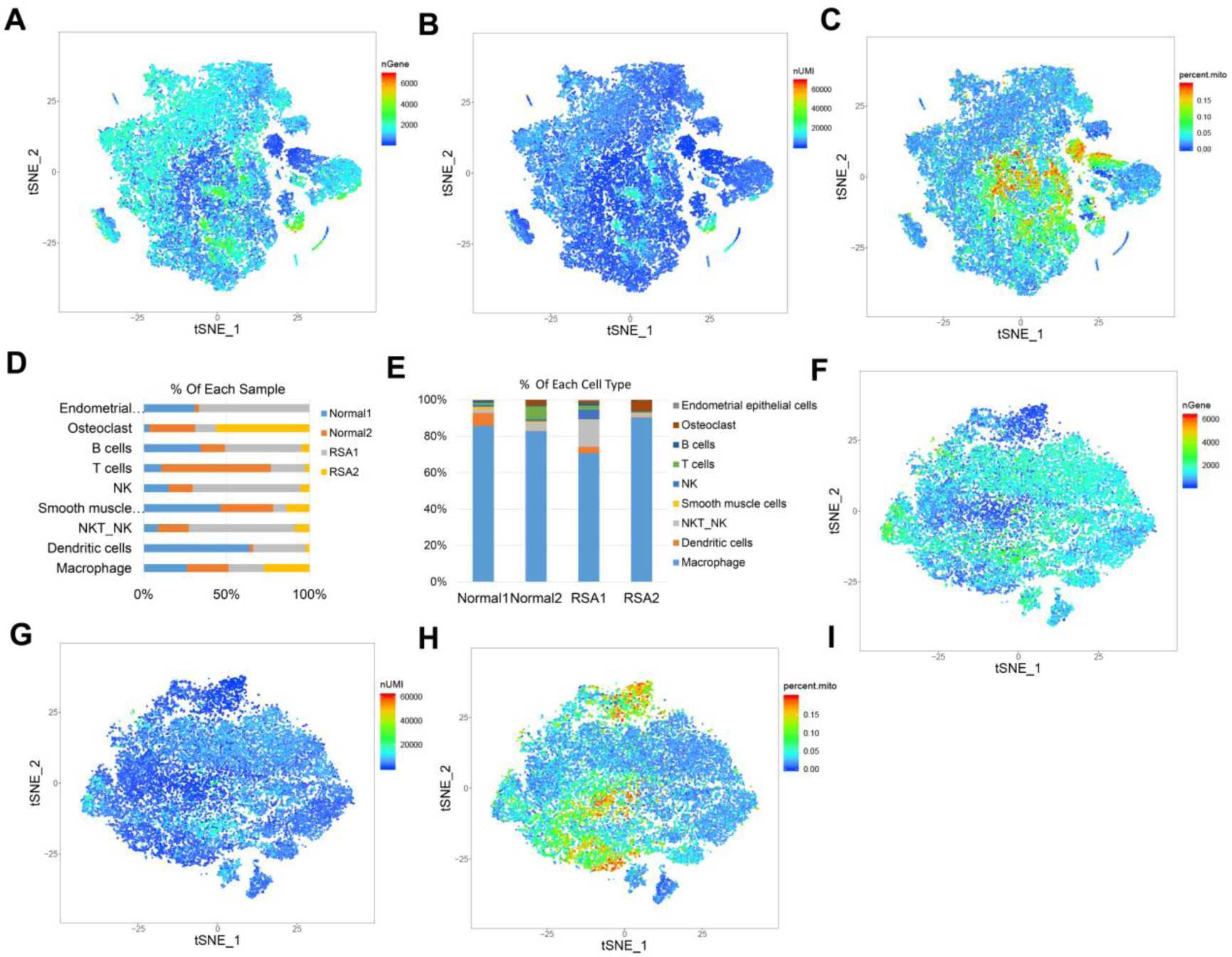
A single-cell analysis of normal and RSA human decidual mononuclear cells. (A-C) t-SNE analysis of decidual mononuclear cells from the combined four samples (Normal 1, Normal 2 and RSA1, RSA2) on nGene (A), nUMI (B) and pecernt mitochondrion (C). (D, E) Percent composition analysis of each cell type in the combined four samples. (F-H) t-SNE analysis of decidual macrophage cells from the combined four samples on nGene (F), nUMI (G) and pecernt mitochondrion (H).

**Figure S4.**
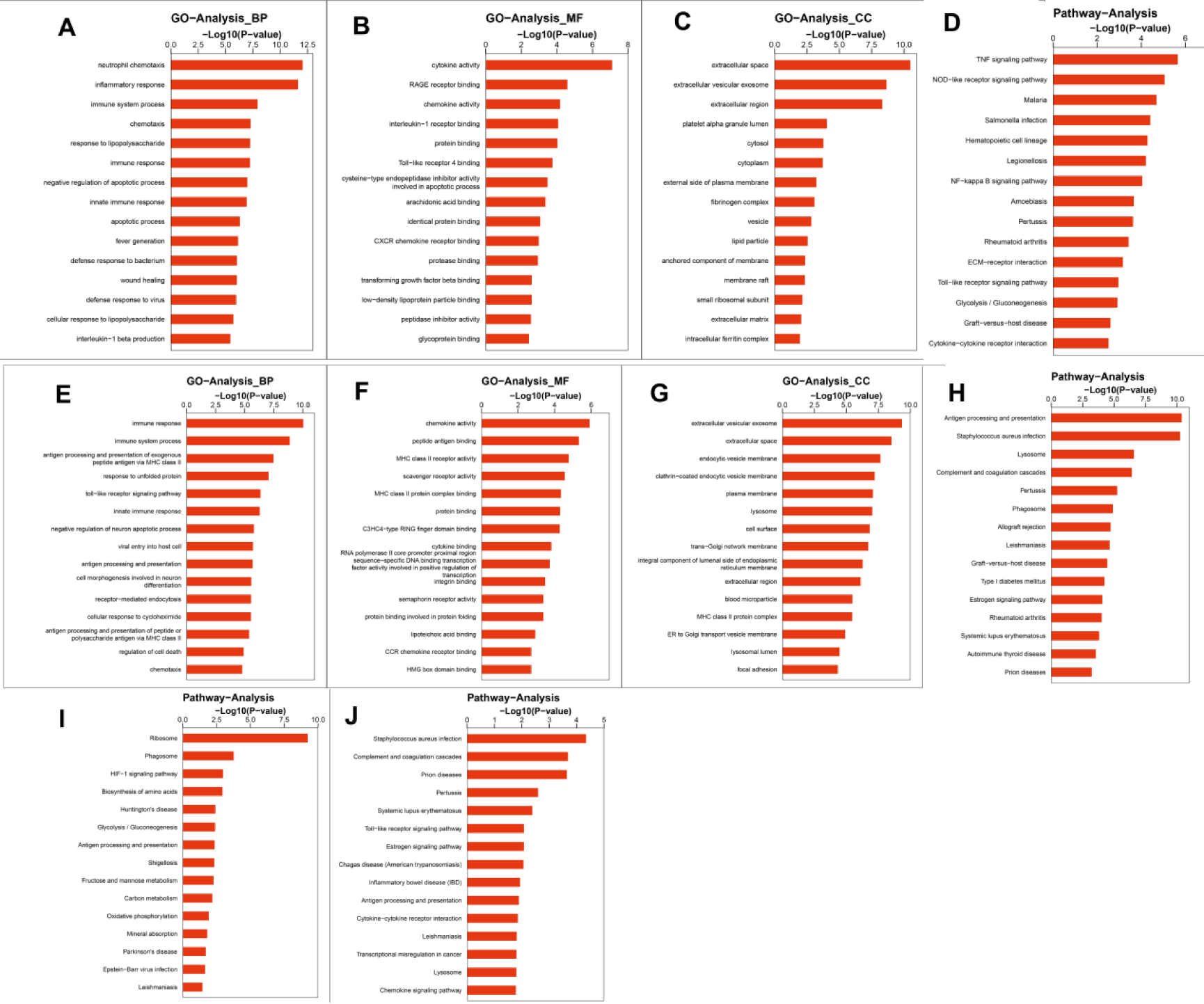
GO and KEGG analysis of macrophage. (A-C) GO analysis of the marker genes in M1 macrophage from normal and RSA sample. (E-G) GO analysis of the marker genes in M2 macrophage from normal and RSA sample. (D,H) KEGG analysis of the marker genes in M1 (D) or M2 (H) macrophage from normal and RSA sample. (I, J) KEGG analysis of the upregulated (I) or downregulated (J) gene in RSA group compared to the normal group.

**Figure S5.**
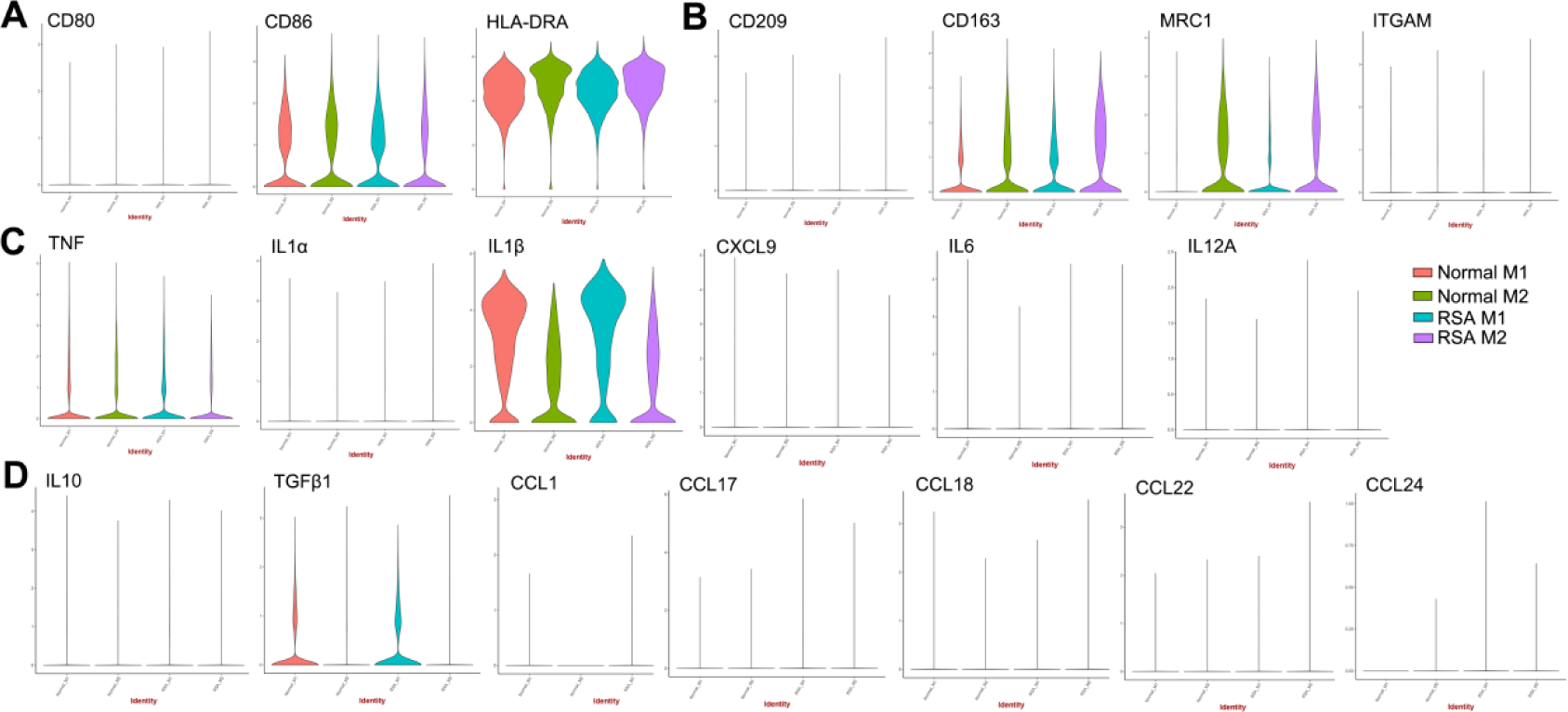
The expression analysis of classically markers and secreted cytokines in the four decidual macrophages. (A,B) Violin plot showing the classically markers of M1 macrophage (A) and M2 macrophage (B). (C,D) Violin plot showing the classically secreted cytokines of M1 macrophage (C) and M2 macrophage (D).

**Figure S6.**
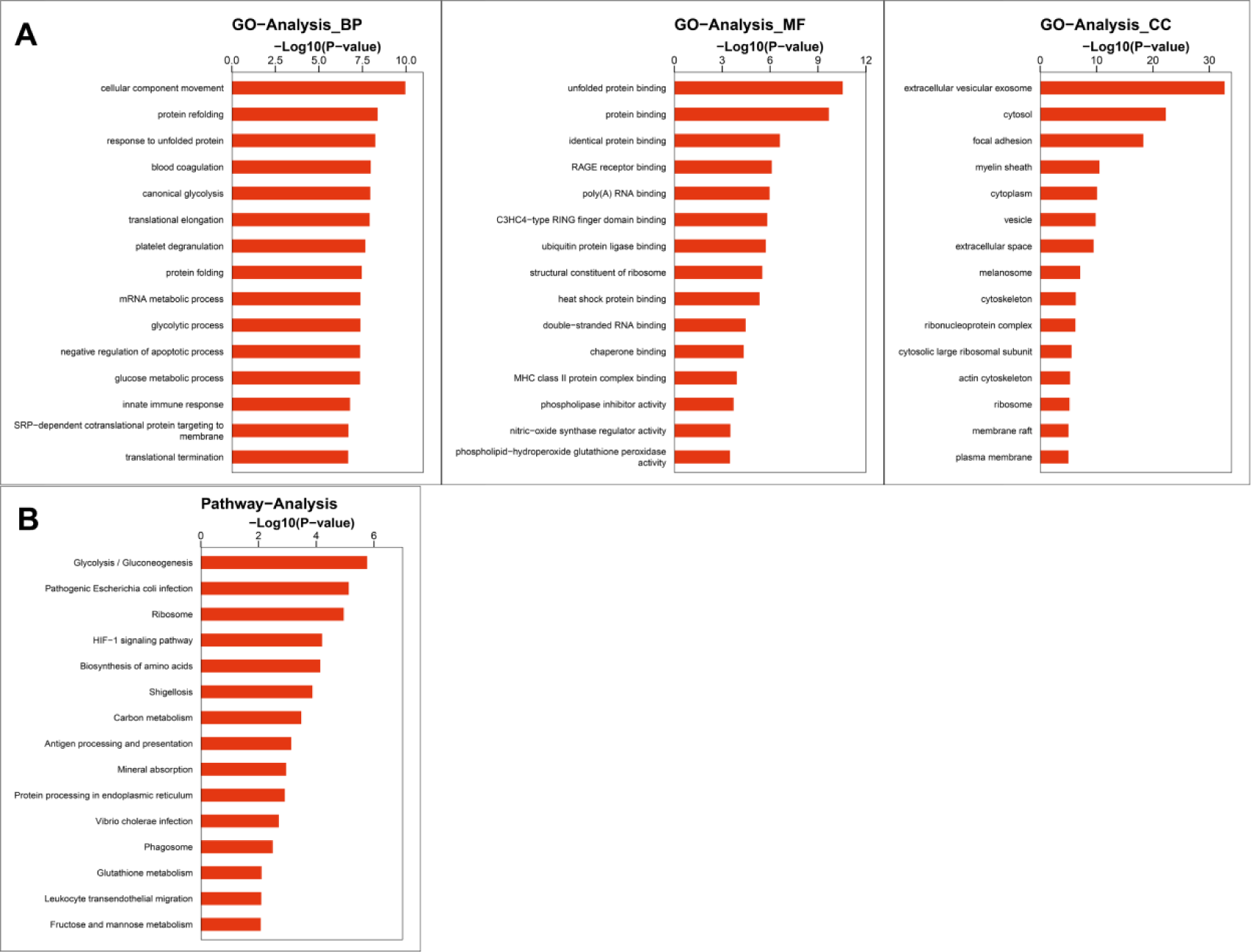
GO and KEGG analysis of Cluster 10 as the specific cluster of decidua macrophage in normal group.

**Figure S7.**
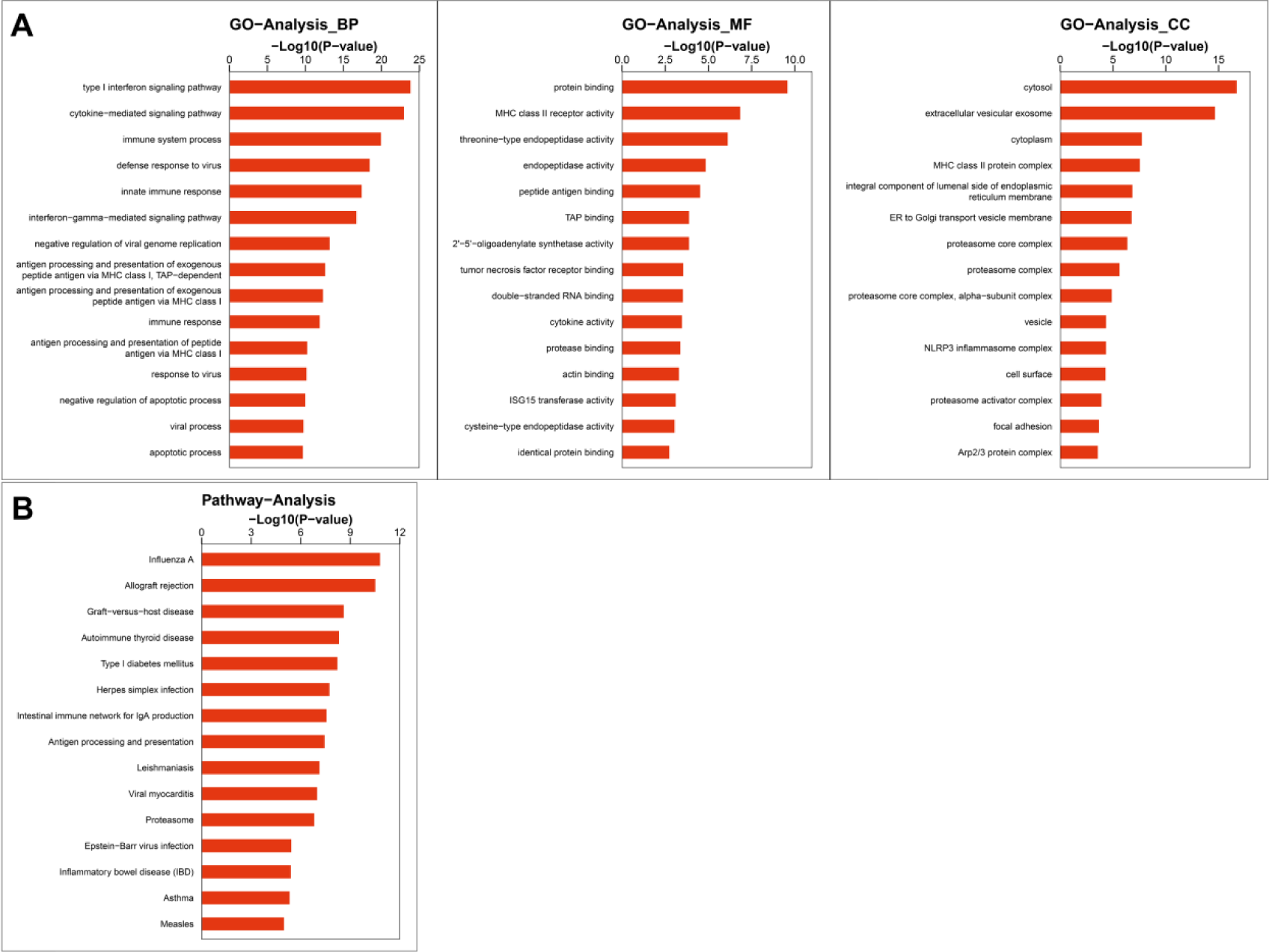
GO and KEGG analysis of Cluster 7 decidua macrophage in normal group.

**Figure S8.**
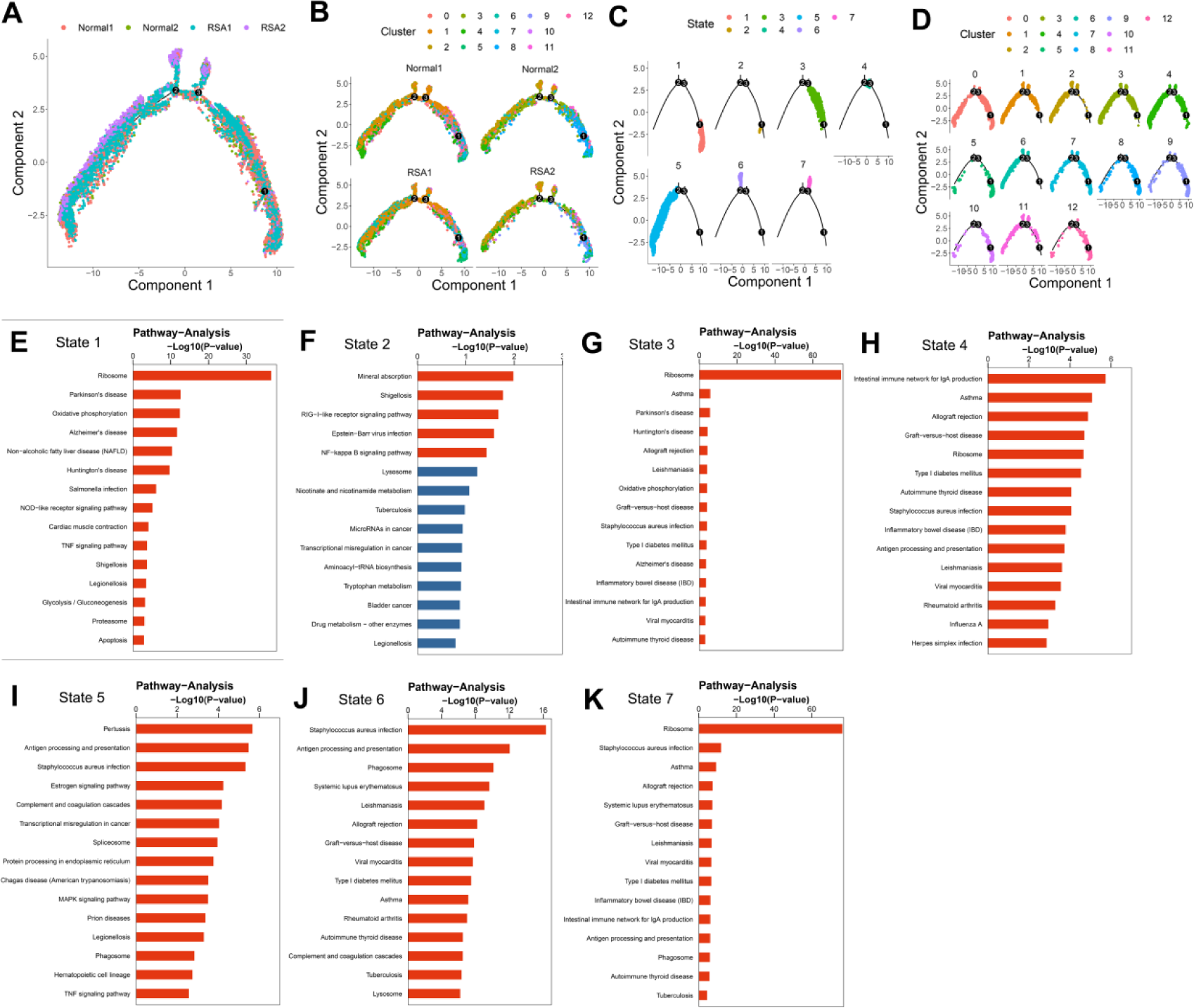
Pseudotime trajectory analysis of decidual macrophage. (A-D) Monocle pseudotime trajectory showing the progression of decidua macrophage. (E-K) KEGG analysis of each state in the pseudotime trajectory analysis.

**Figure S9.**
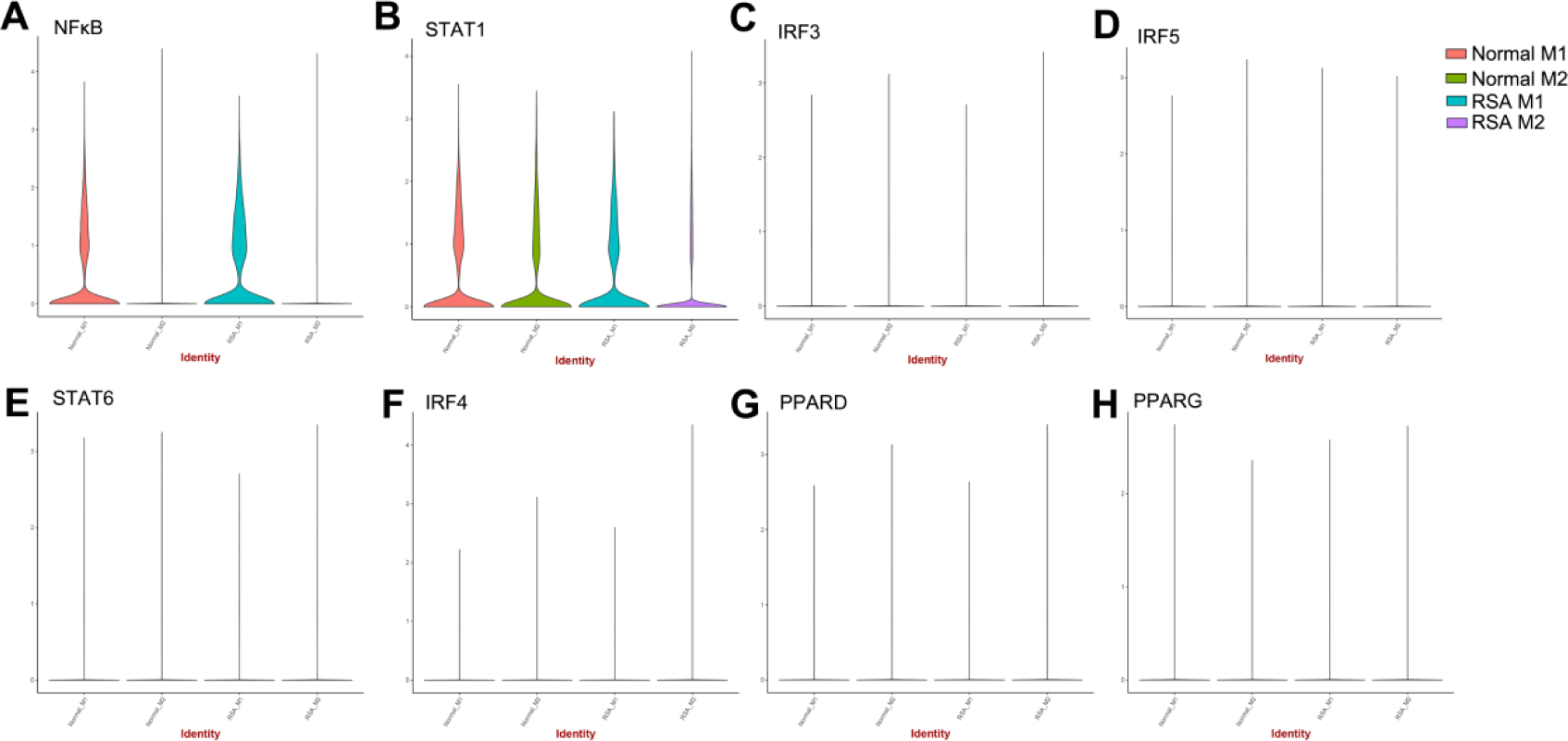
The expression of classically transcription factors in decidua macrophage samples. (A-D) Violin plot showing the expression of transcription factors in M1 type decidua macrophage. (E-H) Violin plot showing the expression of transcription factors in M2 type decidua macrophage.

**Table S1.** Cluster analysis of decidual mononuclear cells in Normal sample

| Cluster | Normal 1 | Normal 2 | Cell Type | Main Markers |
| --- | --- | --- | --- | --- |
| 0 | 1705 | 41 | Macrophage | PTPRC、CD14、MS4A7、CTSS |
| 1 | 2750 | 53 | Macrophage | PTPRC、CD14、MS4A7、CTSS |
| 2 | 736 | 233 | Macrophage | PTPRC、CD14、MS4A7、CTSS |
| 3 | 301 | 1587 | Macrophage | PTPRC、CD14、MS4A7、CTSS |
| 4 | 1372 | 491 | Macrophage | PTPRC、CD14、MS4A7、CTSS |
| 5 | 183 | 794 | Macrophage | PTPRC、CD14、MS4A7、CTSS |
| 6 | 211 | 183 | NKT_NK | PTPRC、NKG7、NCAM1、CD3E、KLRD1 |
| 7 | 744 | 57 | Macrophage | PTPRC、CD14、MS4A7、CTSS |
| 8 | 419 | 81 | Macrophage | PTPRC、CD14、MS4A7、CTSS |
| 9 | 749 | 10 | DC | PTPRC、CD1C |
| 10 | 144 | 210 | Osteoclast | PTPRC、ACP5、RANK、CTSK |
| 11 | 689 | 46 | Macrophage | PTPRC、CD14、MS4A7、CTSS |
| 12 | 126 | 48 | NK | PTPRC、NKG7、NCAM1、KLRD1 |
| 13 | 121 | 299 | T Cells | PTPRC、CD3E、CD8A |
| 14 | 93 | 16 | B&Plasma Cell | PTPRC、CD79A、MS4A1 |
| 15 | 139 | 38 | Smooth muscle cells | TAGLN、ACTA2、DCN、COL1A1 |
| 16 | 31 | 1 | Endometrial epithelial cells | PAEP、GPX3、SLPI |

**Table S2.** M1/M2 analysis of decidua macrophage in Normal sample

| Cluster | Normal 1 | Normal 2 | SubType |
| --- | --- | --- | --- |
| 0 | 2995 | 51 | M1 |
| 1 | 1547 | 86 | M2 |
| 2 | 598 | 478 | M2 |
| 3 | 88 | 1743 | M1 |
| 4 | 487 | 198 | M2 |
| 5 | 1320 | 43 | M1 |
| 6 | 320 | 29 | M2 |
| 7 | 477 | 39 | M1 |
| 8 | 69 | 644 | M1 |
| 9 | 271 | 40 | M1 |
| 10 | 464 | 9 | M1 |
| 11 | 92 | 78 | M2 |
| 12 | 277 | 27 | M1 |

**Table S3.** Cluster analysis of decidual mononuclear cells in Normal and RSA sample

| Cluster | Normal 1 | Normal 2 | RSA 1 | RSA 2 | Cell Type | Main Marker |
| --- | --- | --- | --- | --- | --- | --- |
| 0 | 1705 | 41 | 1953 | 91 | Macrophage | PTPRC、CD14、MS4A7、CTSS |
| 1 | 2750 | 53 | 817 | 65 | Macrophage | PTPRC、CD14、MS4A7、CTSS |
| 2 | 736 | 233 | 1406 | 840 | Macrophage | PTPRC、CD14、MS4A7、CTSS |
| 3 | 301 | 1587 | 386 | 818 | Macrophage | PTPRC、CD14、MS4A7、CTSS |
| 4 | 1372 | 491 | 641 | 114 | Macrophage | PTPRC、CD14、MS4A7、CTSS |
| 5 | 183 | 794 | 409 | 1211 | Macrophage | PTPRC、CD14、MS4A7、CTSS |
| 6 | 211 | 183 | 1477 | 83 | NKT_NK | PTPRC、NKG7、NCAM1、CD3E、KLRD1 |
| 7 | 744 | 57 | 466 | 26 | Macrophage | PTPRC、CD14、MS4A7、CTSS |
| 8 | 419 | 81 | 593 | 170 | Macrophage | PTPRC、CD14、MS4A7、CTSS |
| 9 | 749 | 10 | 348 | 11 | DC | PTPRC、CD1C |
| 10 | 144 | 210 | 350 | 358 | Osteoclast | PTPRC、ACP5、RANK、CTSK |
| 11 | 689 | 46 | 107 | 153 | Macrophage | PTPRC、CD14、MS4A7、CTSS |
| 12 | 126 | 48 | 513 | 17 | NK | PTPRC、NKG7、NCAM1、KLRD1 |
| 13 | 121 | 299 | 220 | 12 | T Cells | PTPRC、CD3E、CD8A |
| 14 | 93 | 16 | 118 | 5 | B&Plasma Cell | PTPRC、CD79A、MS4A1 |
| 15 | 139 | 38 | 22 | 16 | Smooth<br>muscle cells | TAGLN、ACTA2、DCN、COL1A1 |
| 16 | 31 | 1 | 63 | 0 | Endometrial<br>epithelial cells | PAEP、GPX3、SLPI |

**Table S4.** M1/M2 analysis of decidua macrophage in Normal and RSA sample

| Cluster | Normal 1 | Normal 2 | RSA 1 | RSA 2 | SubType |
| --- | --- | --- | --- | --- | --- |
| 0 | 2995 | 51 | 864 | 78 | M1 |
| 1 | 1547 | 86 | 1831 | 75 | M2 |
| 2 | 598 | 478 | 259 | 1701 | M2 |
| 3 | 88 | 1743 | 146 | 655 | M1 |
| 4 | 487 | 198 | 857 | 564 | M1 |
| 5 | 1320 | 43 | 655 | 46 | M1 |
| 6 | 320 | 29 | 1543 | 46 | M2 |
| 7 | 477 | 39 | 365 | 20 | M1 |
| 8 | 69 | 644 | 18 | 107 | M1 |
| 9 | 271 | 40 | 114 | 173 | M1 |
| 10 | 464 | 9 | 21 | 19 | M1 |
| 11 | 92 | 78 | 232 | 100 | M2 |
| 12 | 277 | 27 | 88 | 15 | M1 |

**Table S5.**
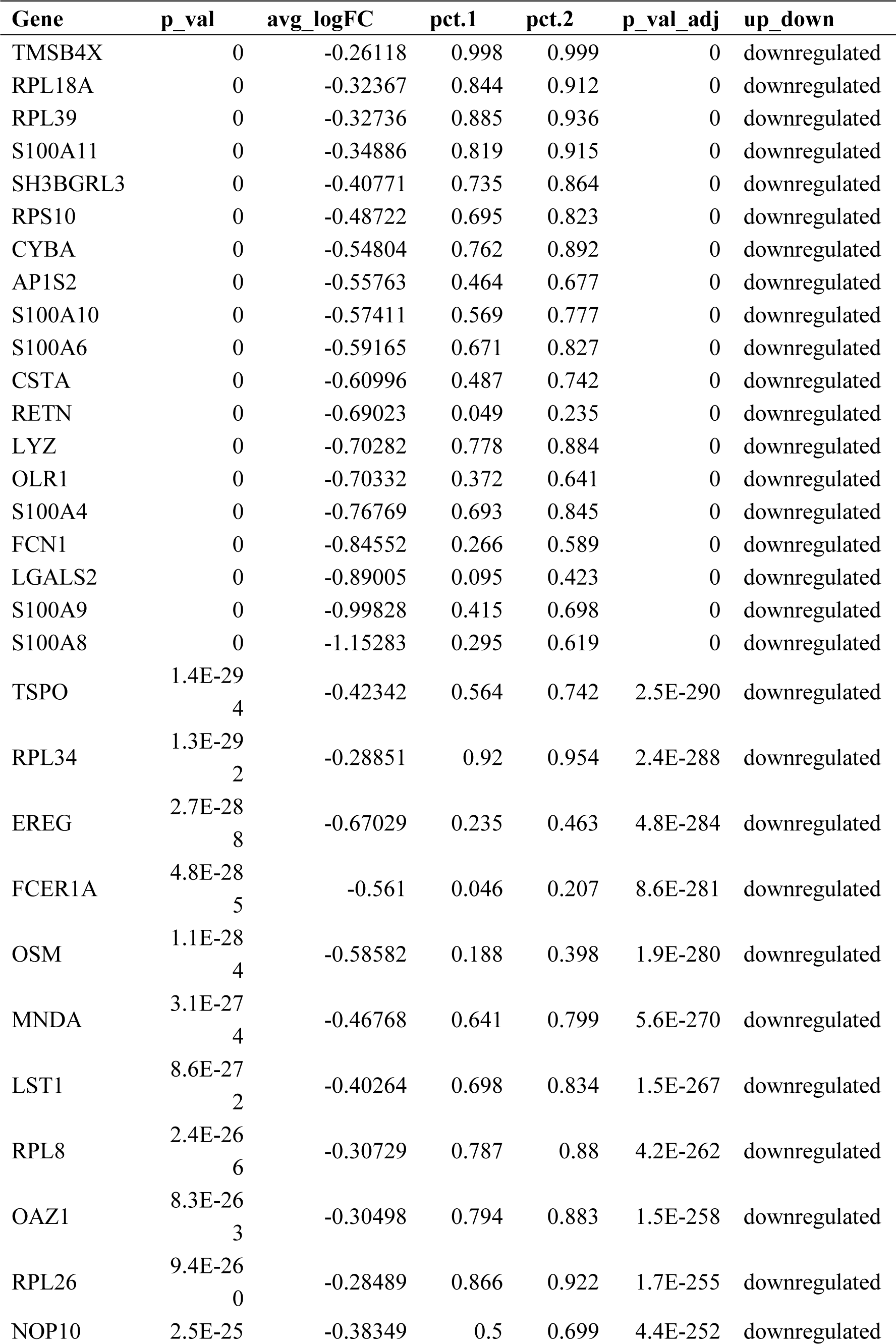
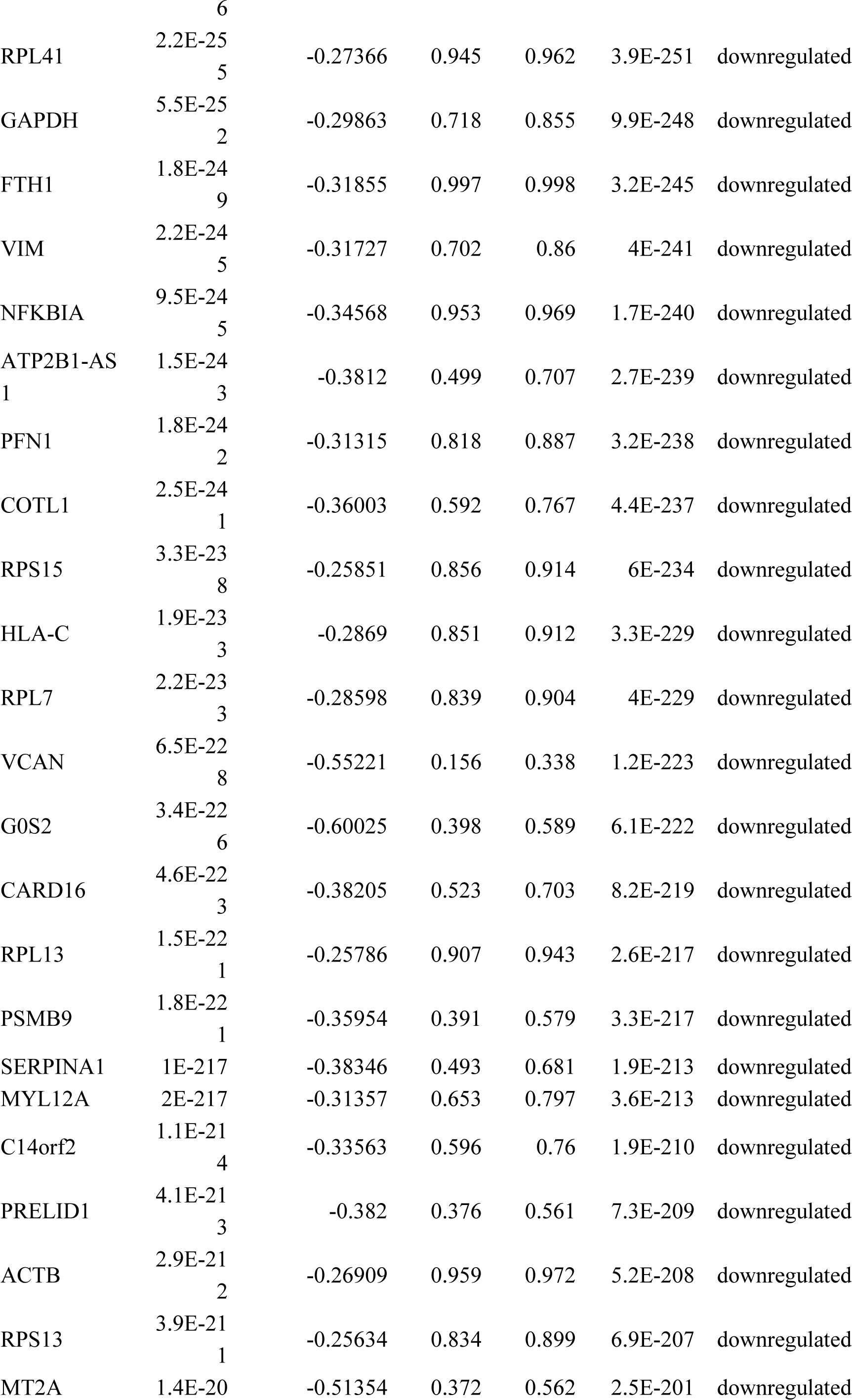
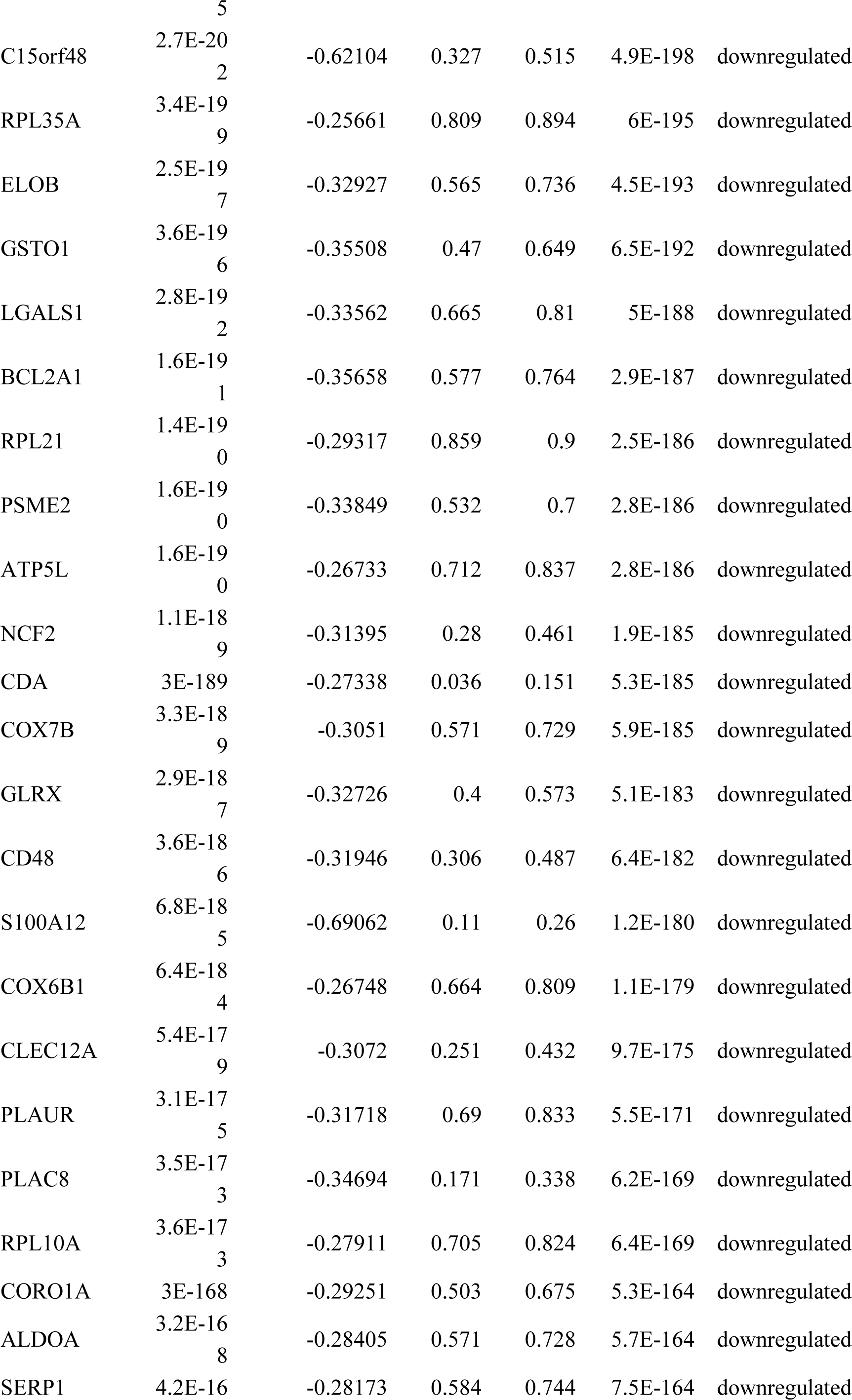
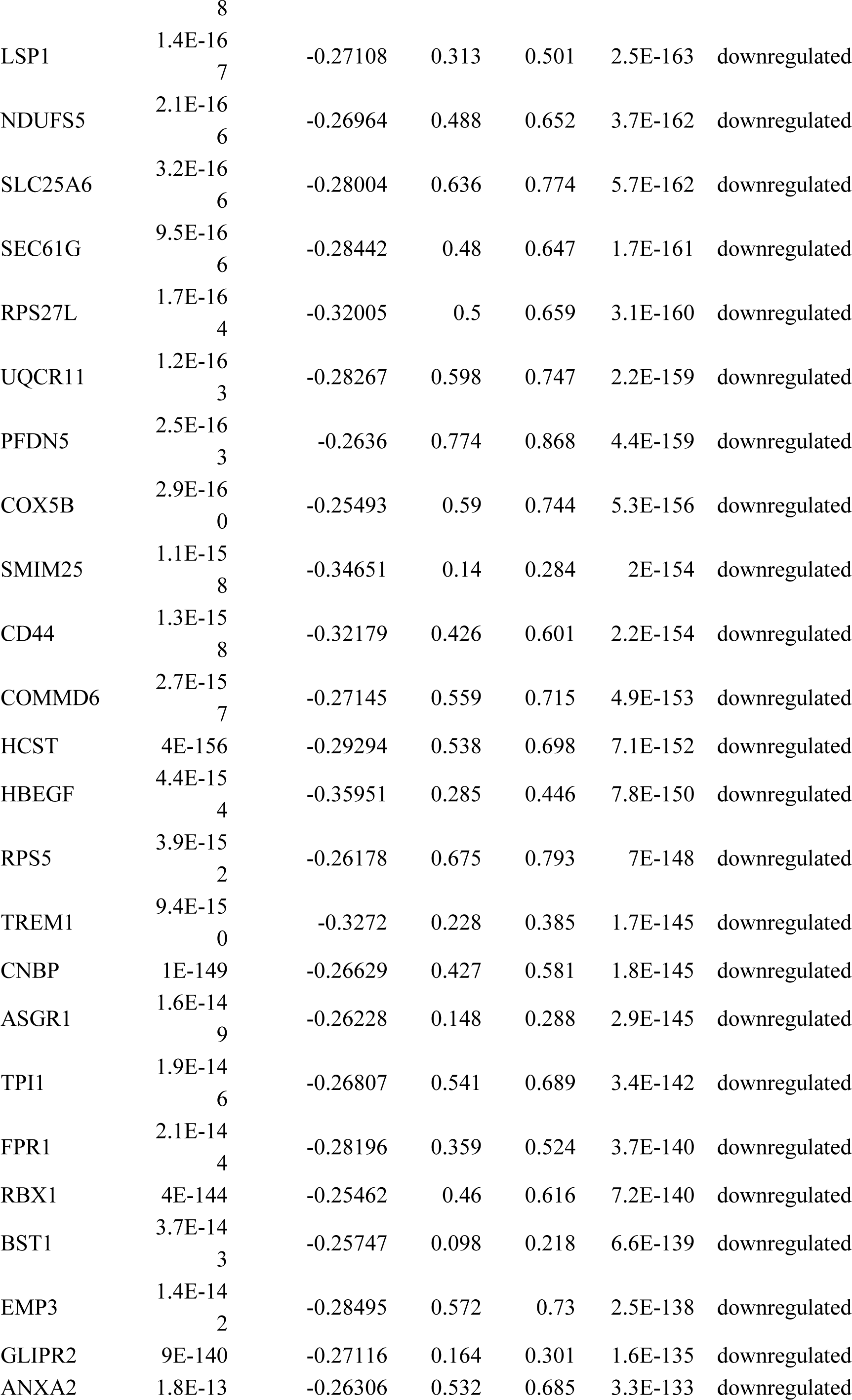
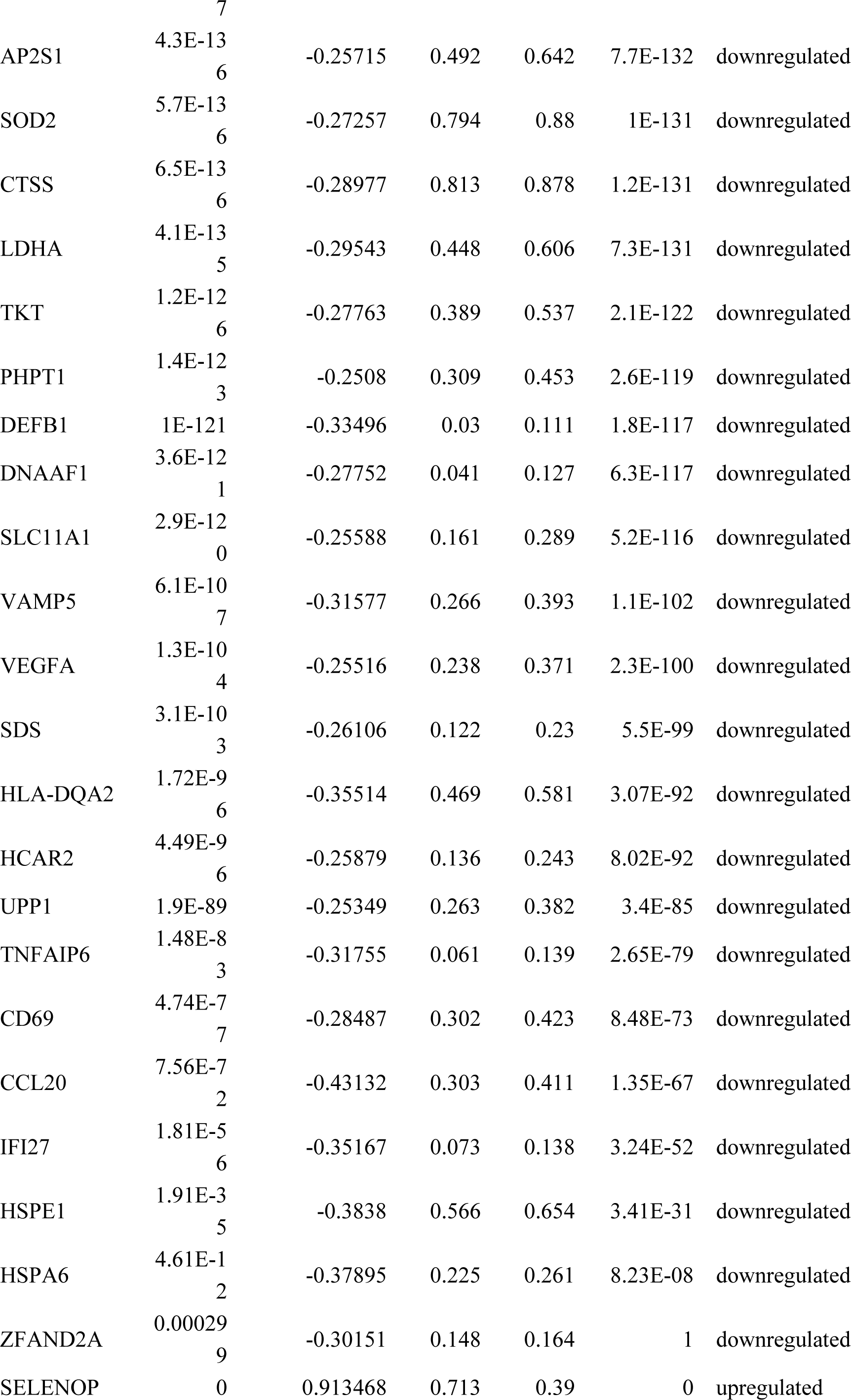
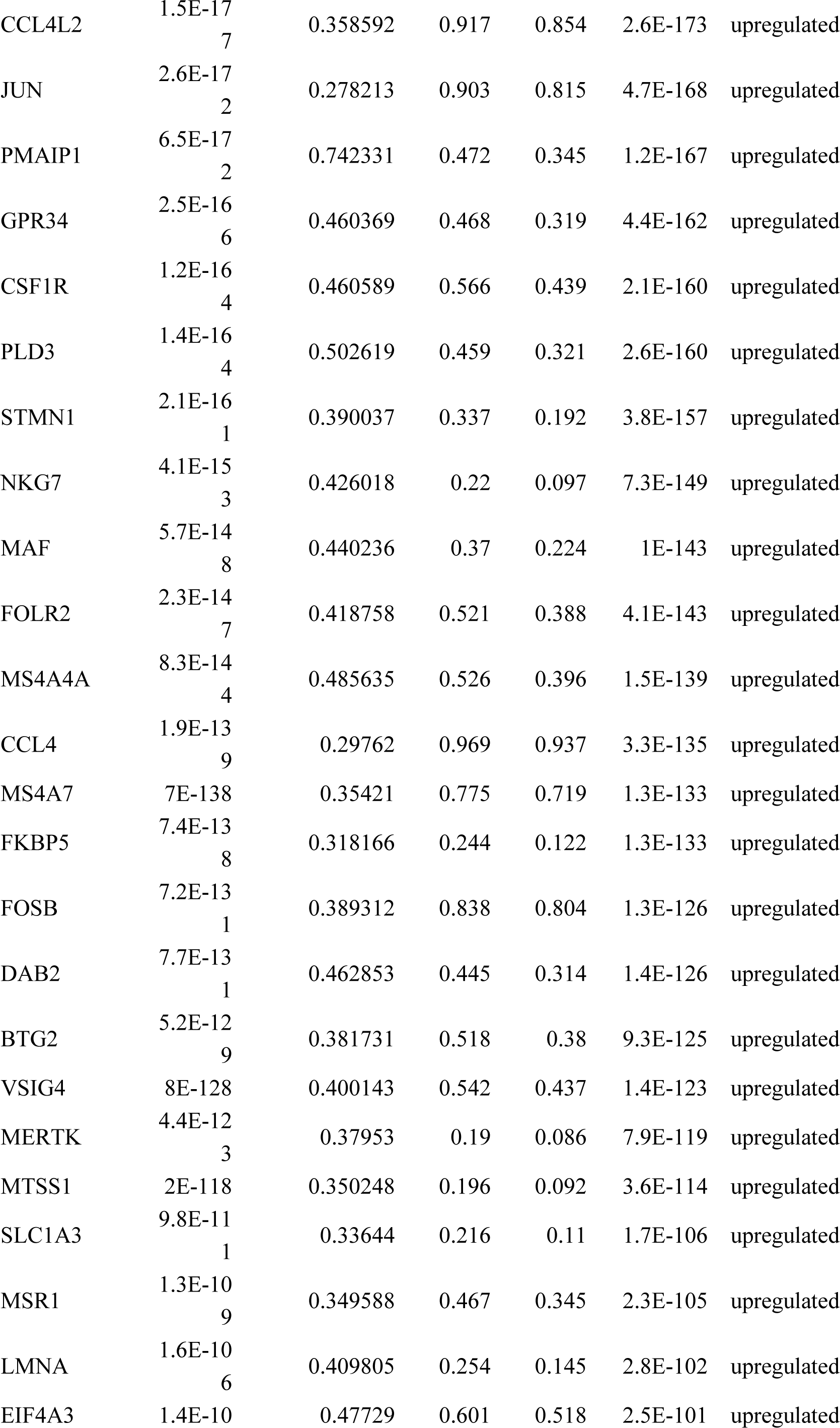
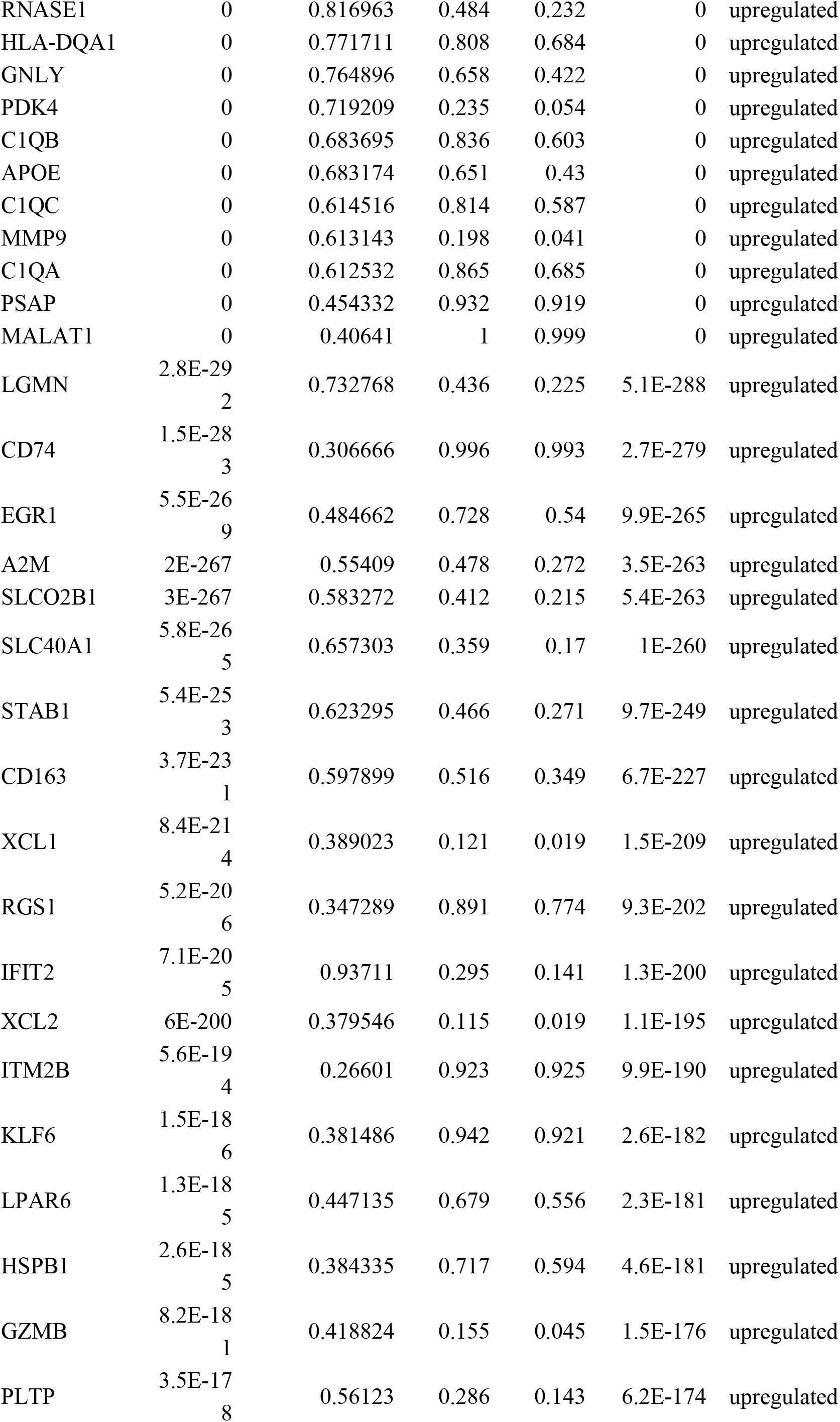
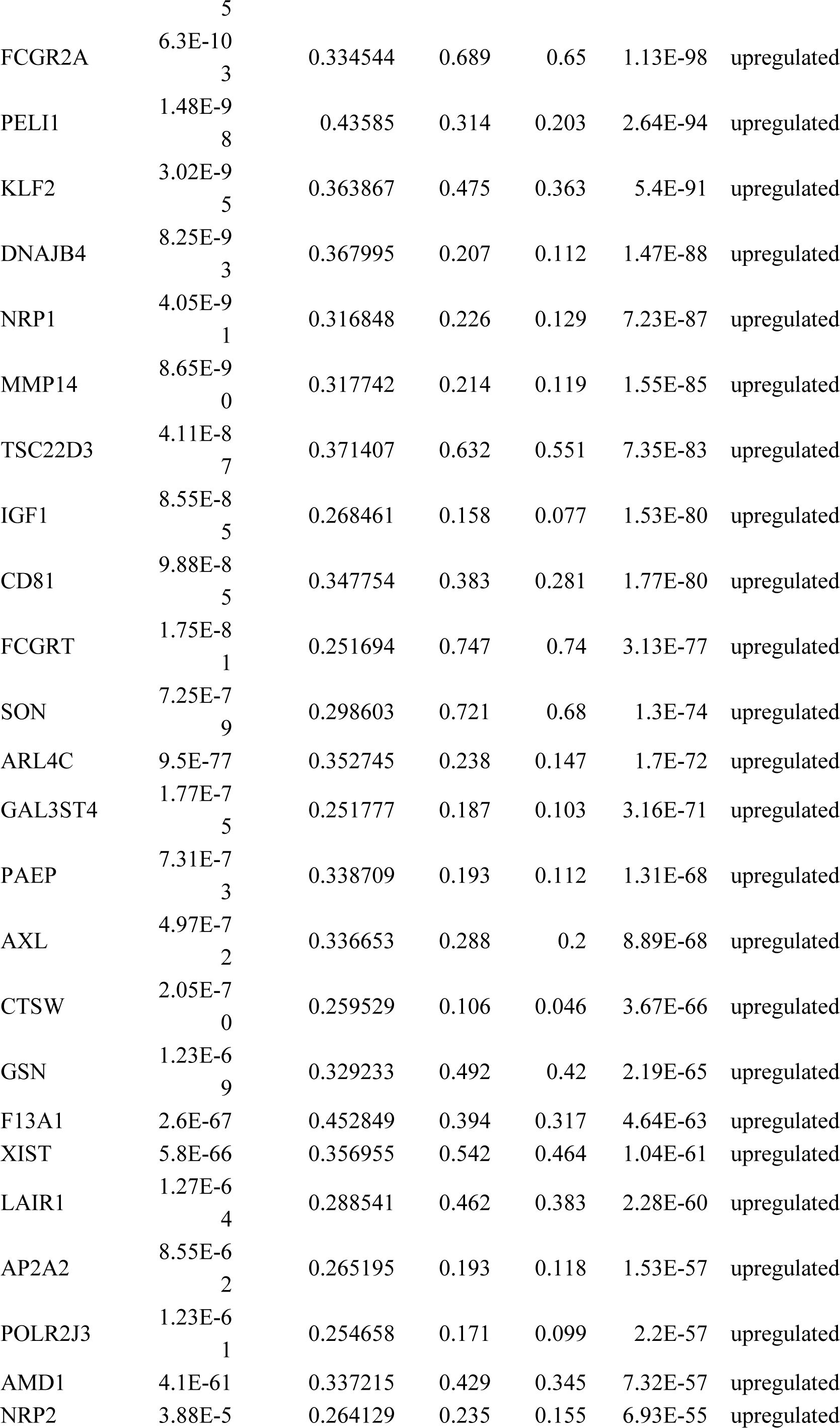
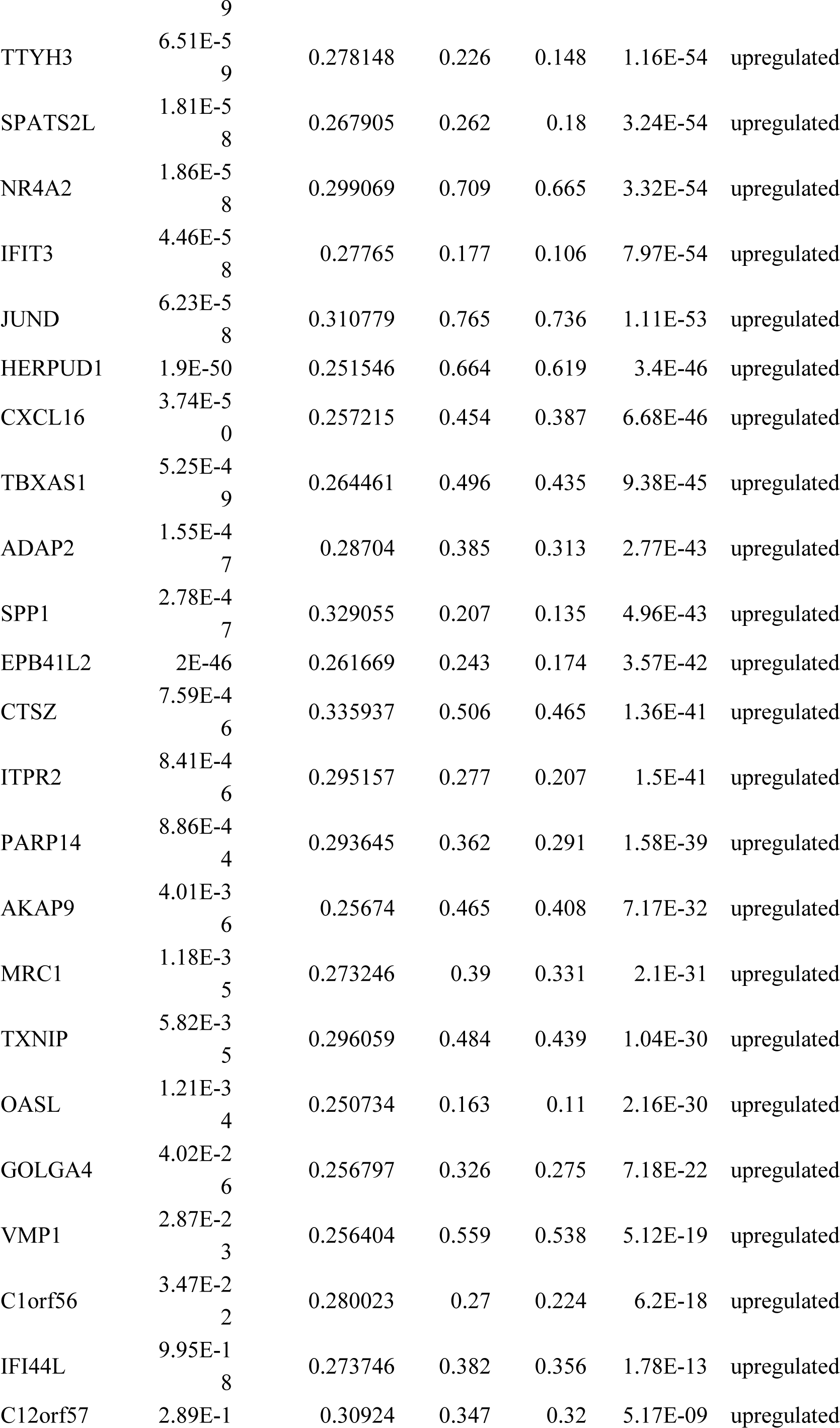
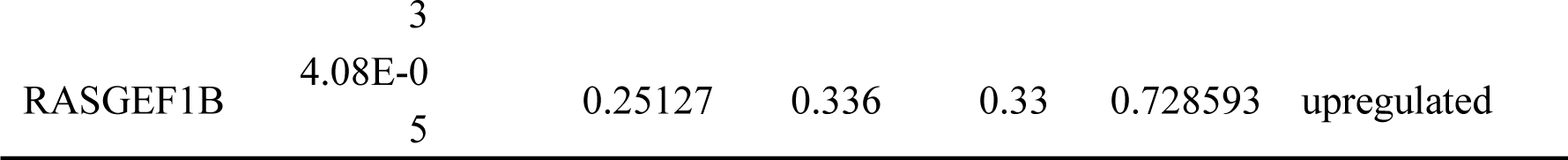
The genes that upregulated (102 genes) and downregulated (122 genes) in the RSA group compared with the normal group.

## REFERENCES

1. Aagaard, K., Ma, J., Antony, K. M., Ganu, R., Petrosino, J., & Versalovic, J. (2014). The placenta harbors a unique microbiome. Sci Transl Med, 6(237), 237–265. doi:10.1126/scitranslmed.3008599

2. Ben Amara, A., Gorvel, L., Baulan, K., Derain-Court, J., Buffat, C., Verollet, C., … Mege, J. L. (2013). Placental macrophages are impaired in chorioamnionitis, an infectious pathology of the placenta. J Immunol, 191(11), 5501–5514. doi:10.4049/jimmunol.1300988

3. Bockle, B. C., Solder, E., Kind, S., Romani, N., & Sepp, N. T. (2008). DC-sign+ CD163+ macrophages expressing hyaluronan receptor LYVE-1 are located within chorion villi of the placenta. Placenta, 29(2), 187–192. doi:10.1016/j.placenta.2007.11.003

4. Brown, M. B., von Chamier, M., Allam, A. B., & Reyes, L. (2014). M1/M2 macrophage polarity in normal and complicated pregnancy. Front Immunol, 5, 606. doi:10.3389/fimmu.2014.00606

5. Buckley, R. J., Whitley, G. S., Dumitriu, I. E., & Cartwright, J. E. (2016). Macrophage polarisation affects their regulation of trophoblast behaviour. Placenta, 47, 73–80. doi:10.1016/j.placenta.2016.09.004

6. Cappelletti, M., Presicce, P., Lawson, M. J., Chaturvedi, V., Stankiewicz, T. E., Vanoni, S., … Divanovic, S. (2017). Type I interferons regulate susceptibility to inflammation-induced preterm birth. JCI Insight, 2(5), e91288. doi:10.1172/jci.insight.91288

7. Deroux, A., Dumestre-Perard, C., Dunand-Faure, C., Bouillet, L., & Hoffmann, P. (2017). Female Infertility and Serum Auto-antibodies: a Systematic Review. Clin Rev Allergy Immunol, 53(1), 78–86. doi:10.1007/s12016-016-8586-z

8. Ding, J., Yang, C., Cheng, Y., Wang, J., Zhang, S., Yan, S., … Yang, J. (2020). Trophoblast-derived IL-6 serves as an important factor for normal pregnancy by activating Stat3-mediated M2 macrophages polarization. Int Immunopharmacol, 106788. doi:10.1016/j.intimp.2020.106788

9. Erlebacher, A. (2013). Immunology of the maternal-fetal interface. Annu Rev Immunol, 31, 387–411. doi:10.1146/annurev-immunol-032712-100003

10. Ferreira, L. M., Meissner, T. B., Tilburgs, T., & Strominger, J. L. (2017). HLA-G: At the Interface of Maternal-Fetal Tolerance. Trends Immunol, 38(4), 272–286. doi:10.1016/j.it.2017.01.009

11. Garcia-Gomez, E., Vazquez-Martinez, E. R., Reyes-Mayoral, C., Cruz-Orozco, O. P., Camacho-Arroyo, I., & Cerbon, M. (2019). Regulation of Inflammation Pathways and Inflammasome by Sex Steroid Hormones in Endometriosis. Front Endocrinol (Lausanne), 10, 935. doi:10.3389/fendo.2019.00935

12. Gomez-Lopez, N., StLouis, D., Lehr, M. A., Sanchez-Rodriguez, E. N., & Arenas-Hernandez, M. (2014). Immune cells in term and preterm labor. Cell Mol Immunol, 11(6), 571–581. doi:10.1038/cmi.2014.46

13. Han, X., Chen, H., Huang, D., Chen, H., Fei, L., Cheng, C., … Guo, G. (2018). Mapping human pluripotent stem cell differentiation pathways using high throughput single-cell RNA-sequencing. Genome Biol, 19(1), 47. doi:10.1186/s13059-018-1426-0

14. Houser, B. L., Tilburgs, T., Hill, J., Nicotra, M. L., & Strominger, J. L. (2011). Two unique human decidual macrophage populations. J Immunol, 186(4), 2633–2642. doi:10.4049/jimmunol.1003153

15. Hu, X. H., Tang, M. X., Mor, G., & Liao, A. H. (2016). Tim-3: Expression on immune cells and roles at the maternal-fetal interface. J Reprod Immunol, 118, 92–99. doi:10.1016/j.jri.2016.10.113

16. Huhn, O., Ivarsson, M. A., Gardner, L., Hollinshead, M., Stinchcombe, J. C., Chen, P., … Colucci, F. (2020). Distinctive phenotypes and functions of innate lymphoid cells in human decidua during early pregnancy. Nat Commun, 11(1), 381. doi:10.1038/s41467-019-14123-z

17. Jena, M. K., Nayak, N., Chen, K., & Nayak, N. R. (2019). Role of Macrophages in Pregnancy and Related Complications. Arch Immunol Ther Exp (Warsz), 67(5), 295–309. doi:10.1007/s00005-019-00552-7

18. Kammerer, U., Eggert, A. O., Kapp, M., McLellan, A. D., Geijtenbeek, T. B., Dietl, J., … Kampgen, E. (2003). Unique appearance of proliferating antigen-presenting cells expressing DC-SIGN (CD209) in the decidua of early human pregnancy. Am J Pathol, 162(3), 887–896. doi:10.1016/S0002-9440(10)63884-9

19. Kang, X., Zhang, X., & Zhao, A. (2016). Macrophage depletion and TNF-alpha inhibition prevent resorption in CBA/J x DBA/2 model of CpG-induced abortion. Biochem Biophys Res Commun, 469(3), 704–710. doi:10.1016/j.bbrc.2015.12.024

20. Khalaj, K., Luna, R. L., de Franca, M. E., de Oliveira, W. H., Peixoto, C. A., & Tayade, C. (2016). RNA binding protein, tristetraprolin in a murine model of recurrent pregnancy loss. Oncotarget, 7(45), 72486–72502. doi:10.18632/oncotarget.12539

21. Li, Y. H., Zhou, W. H., Tao, Y., Wang, S. C., Jiang, Y. L., Zhang, D., … Du, M. R. (2016). The Galectin-9/Tim-3 pathway is involved in the regulation of NK cell function at the maternal-fetal interface in early pregnancy. Cell Mol Immunol, 13(1), 73–81. doi:10.1038/cmi.2014.126

22. Liu, S., Diao, L., Huang, C., Li, Y., Zeng, Y., & Kwak-Kim, J. Y. H. (2017). The role of decidual immune cells on human pregnancy. J Reprod Immunol, 124, 44–53. doi:10.1016/j.jri.2017.10.045

23. Macklin, P. S., McAuliffe, J., Pugh, C. W., & Yamamoto, A. (2017). Hypoxia and HIF pathway in cancer and the placenta. Placenta, 56, 8–13. doi:10.1016/j.placenta.2017.03.010

24. Ning, F., Liu, H., & Lash, G. E. (2016). The Role of Decidual Macrophages During Normal and Pathological Pregnancy. Am J Reprod Immunol, 75(3), 298–309. doi:10.1111/aji.12477

25. Olmos-Ortiz, A., Flores-Espinosa, P., Mancilla-Herrera, I., Vega-Sanchez, R., Diaz, L., & Zaga-Clavellina, V. (2019). Innate Immune Cells and Toll-like Receptor-Dependent Responses at the Maternal-Fetal Interface. Int J Mol Sci, 20(15), 3654. doi:10.3390/ijms20153654

26. Ono, Y., Nagai, M., Yoshino, O., Koga, K., Nawaz, A., Hatta, H., … Saito, S. (2018). CD11c+ M1-like macrophages (MPhis) but not CD206+ M2-like MPhi are involved in folliculogenesis in mice ovary. Sci Rep, 8(1), 8171. doi:10.1038/s41598-018-25837-3

27. Perez-Sepulveda, A., Torres, M. J., Khoury, M., & Illanes, S. E. (2014). Innate immune system and preeclampsia. Front Immunol, 5, 244. doi:10.3389/fimmu.2014.00244

28. Picelli, S., Faridani, O. R., Bjorklund, A. K., Winberg, G., Sagasser, S., & Sandberg, R. (2014). Full-length RNA-seq from single cells using Smart-seq2. Nat Protoc, 9(1), 171–181. doi:10.1038/nprot.2014.006

29. PrabhuDas, M., Bonney, E., Caron, K., Dey, S., Erlebacher, A., Fazleabas, A., … Yoshinaga, K. (2015). Immune mechanisms at the maternal-fetal interface: perspectives and challenges. Nat Immunol, 16(4), 328–334. doi:10.1038/ni.3131

30. Raj, R. S., Bonney, E. A., & Phillippe, M. (2014). Influenza, immune system, and pregnancy. Reprod Sci, 21(12), 1434–1451. doi:10.1177/1933719114537720

31. Rosenberg, A. B., Roco, C. M., Muscat, R. A., Kuchina, A., Sample, P., Yao, Z., … Seelig, G. (2018). Single-cell profiling of the developing mouse brain and spinal cord with split-pool barcoding. Science, 360(6385), 176–182. doi:10.1126/science.aam8999

32. Soares, M. J., Iqbal, K., & Kozai, K. (2017). Hypoxia and Placental Development. Birth Defects Res, 109(17), 1309–1329. doi:10.1002/bdr2.1135

33. Sun, H., Wen, X., Li, H., Wu, P., Gu, M., Zhao, X., … Zhang, Z. (2020). Single-cell RNA-seq analysis identifies meniscus progenitors and reveals the progression of meniscus degeneration. Ann Rheum Dis, 79(3), 408–417. doi:10.1136/annrheumdis-2019-215926

34. Svensson-Arvelund, J., & Ernerudh, J. (2015). The Role of Macrophages in Promoting and Maintaining Homeostasis at the Fetal-Maternal Interface. Am J Reprod Immunol, 74(2), 100–109. doi:10.1111/aji.12357

35. Svensson, J., Jenmalm, M. C., Matussek, A., Geffers, R., Berg, G., & Ernerudh, J. (2011). Macrophages at the fetal-maternal interface express markers of alternative activation and are induced by M-CSF and IL-10. J Immunol, 187(7), 3671–3682. doi:10.4049/jimmunol.1100130

36. Ticconi, C., Pietropolli, A., Di Simone, N., Piccione, E., & Fazleabas, A. (2019). Endometrial Immune Dysfunction in Recurrent Pregnancy Loss. Int J Mol Sci, 20(21), 5332. doi:10.3390/ijms20215332

37. Tilburgs, T., Evans, J. H., Crespo, A. C., & Strominger, J. L. (2015). The HLA-G cycle provides for both NK tolerance and immunity at the maternal-fetal interface. Proc Natl Acad Sci U S A, 112(43), 13312–13317. doi:10.1073/pnas.1517724112

38. Triggianese, P., Perricone, C., Chimenti, M. S., De Carolis, C., & Perricone, R. (2016). Innate Immune System at the Maternal-Fetal Interface: Mechanisms of Disease and Targets of Therapy in Pregnancy Syndromes. Am J Reprod Immunol, 76(4), 245–257. doi:10.1111/aji.12509

39. Tsang, J. C. H., Vong, J. S. L., Ji, L., Poon, L. C. Y., Jiang, P., Lui, K. O., … Lo, Y. M. D. (2017). Integrative single-cell and cell-free plasma RNA transcriptomics elucidates placental cellular dynamics. Proc Natl Acad Sci U S A, 114(37), E7786–E7795. doi:10.1073/pnas.1710470114

40. Tsao, F. Y., Wu, M. Y., Chang, Y. L., Wu, C. T., & Ho, H. N. (2018). M1 macrophages decrease in the deciduae from normal pregnancies but not from spontaneous abortions or unexplained recurrent spontaneous abortions. J Formos Med Assoc, 117(3), 204–211. doi:10.1016/j.jfma.2017.03.011

41. Vento-Tormo, R., Efremova, M., Botting, R. A., Turco, M. Y., Vento-Tormo, M., Meyer, K. B., … Teichmann, S. A. (2018). Single-cell reconstruction of the early maternal-fetal interface in humans. Nature, 563(7731), 347–353. doi:10.1038/s41586-018-0698-6

42. Vishnyakova, P., Elchaninov, A., Fatkhudinov, T., & Sukhikh, G. (2019). Role of the Monocyte-Macrophage System in Normal Pregnancy and Preeclampsia. Int J Mol Sci, 20(15), 3695. doi:10.3390/ijms20153695

43. Yang, F., Zheng, Q., & Jin, L. (2019). Dynamic Function and Composition Changes of Immune Cells During Normal and Pathological Pregnancy at the Maternal-Fetal Interface. Front Immunol, 10, 2317. doi:10.3389/fimmu.2019.02317

44. Zhang, Y. H., He, M., Wang, Y., & Liao, A. H. (2017). Modulators of the Balance between M1 and M2 Macrophages during Pregnancy. Front Immunol, 8, 120. doi:10.3389/fimmu.2017.00120

45. Zheng, Q., Hou, J., Zhou, Y., Li, Z., & Cao, X. (2017). The RNA helicase DDX46 inhibits innate immunity by entrapping m(6)A-demethylated antiviral transcripts in the nucleus. Nat Immunol, 18(10), 1094–1103. doi:10.1038/ni.3830

